# SenSet, a novel human lung senescence cell gene signature, identifies cell-specific senescence mechanisms

**DOI:** 10.1101/2024.12.21.629928

**Authors:** Euxhen Hasanaj, Delphine Beaulieu, Cankun Wang, Qianjiang Hu, Marta Bueno, John C Sembrat, Ricardo H Pineda, Maria Camila Melo-Narvaez, Nayra Cardenes, Zhao Yanwu, Zhang Yingze, Robert Lafyatis, Alison Morris, Ana Mora, Mauricio Rojas, Dongmei Li, Irfan Rahman, Gloria S Pryhuber, Mareike Lehmann, Jonathan Alder, Aditi Gurkar, Toren Finkel, Qin Ma, Barnabás Póczos, Ziv Bar-Joseph, Oliver Eickelberg, Melanie Königshoff

## Abstract

Cellular senescence is a major hallmark of aging. Senescence is defined as an irreversible growth arrest observed when cells are exposed to a variety of stressors including DNA damage, oxidative stress, or nutrient deprivation. While senescence is a well-established driver of aging and age-related diseases, it is a highly heterogeneous process with significant variations across organisms, tissues, and cell types.

The relatively low abundance of senescence in healthy aged tissues represents a major challenge to studying senescence in a given organ, including the human lung. To overcome this limitation, we developed a Positive-Unlabeled (PU) learning framework to generate a comprehensive senescence marker gene list in human lungs (termed SenSet) using the largest publicly available single-cell lung dataset, the Human Lung Cell Atlas (HLCA). We validated SenSet in a highly complex ex vivo human 3D lung tissue culture model subjected to the senescence inducers bleomycin, doxorubicin, or irradiation, and established its sensitivity and accuracy in characterizing senescence. Using SenSet, we identified and validated cell-type specific senescence signatures in distinct lung cell populations upon aging and environmental exposures. Our study presents the first comprehensive analysis of senescent cells in the healthy aging lung and uncovers cell-specific gene signatures of senescence, presenting fundamental implications for our understanding of major lung diseases, including cancer, fibrosis, chronic obstructive pulmonary disease, or asthma.

## Introduction

Cellular senescence refers to a permanent arrest of cell division, triggered by a variety of stressors or insults. DNA damage is a common inducer of senescence, which can occur over time or in response to oxidative stress, nutrient deprivation, and oncogenic signaling, among others^1–6^. The absence of cell division can detrimentally impact tissue regeneration and repair, thereby contributing to numerous age-related diseases, including but not limited to cardiopulmonary and neurodegenerative diseases or metabolic syndromes^7–9^. Senescent cells (SnCs) are resistant to apoptosis, which at least partially accounts for the accumulation of SnCs in advanced age. Moreover, SnCs have been known to undergo the senescence associated secretory phenotype (SASP), further exacerbating tissue dysfunction in aged individuals by selective secretion of soluble mediators driving senescence in adjacent cells^10^. Both SASP and senescence markers are highly dependent on cell type^11^, as such, the precise characterization of SnC identity, especially in healthy tissues over age, remains imperative for understanding aging. Hence, we pursue a more refined definition of cell-type specific senescence markers in this contribution.

Discovery of genetic markers for senescence is a first and crucial step in understanding the mechanisms involved. Not only will such markers facilitate the identification of SnCs, but they will also allow us to track the trajectory of SnC development, their spatial location, and the cell-cell interactions that drive senescence. As such, senescence markers could help identify potential targets for therapeutic interventions, particularly when removal of SnCs could be beneficial.

A number of prior studies and methods attempted to compile marker gene sets enriched for senescence based on gene knockout models, functional assays, and literature reviews. For instance, SenMayo is a gene set of 125 genes that were reported to be upregulated in human or mouse senescent cells^12^. Similarly, Fridman & Tainsky enumerate several genes that are involved in senescence-related pathways^13^, and CellAge is another curated database of 279 genes driving senescence^14^. A limitation of using these sets is their limited overlap and the lack of validation in an independent, large cohort of aging individuals.

In this work, we sought to identify a refined list of marker genes with high sensitivity for senescence in human lungs by drawing upon these prior gene sets. Although the genes in these prior sets may not be universally consistent, through the lens of information theory, they can be considered as partial or noisy information for labeling senescence. For this, we extended methods for *weakly supervised learning* which are aimed at effectively leveraging incomplete information to classify samples^15^ (in our case, to determine if a cell is senescent or not). The importance of such a weakly supervised approach lies in its ability to handle the inherent variability and complexity of biological systems, where a gene may have multiple functions, including roles beyond the promotion of senescence. Specifically, we used a class of learners known as positive-unlabeled (PU) learning algorithms^16^ to identify SnCs. In our case, we applied PU learning to a large cohort of single-cell lung data from individuals encompassing the human life span. While it is currently impossible to definitively identify which cells in this cohort are senescent, we use PU learning to isolate a subset of cells from older individuals that differ from healthy young cells, restricted to genes associated with senescence as reported in the literature.

Based on our PU learning approach, we identified a subset of genes from four combined prior senescence marker lists: GO:0090398, Fridman, SenMayo, and CellAge^12–14,17^ that showed significant senescence-related activity. The genes were selected by performing PU learning using different age groups in the Human Lung Cell Atlas (HLCA)^18^, which combines data from 106 healthy donors spanning an age range from 10 to 76. We termed the resulting list SenSet. Next, we extensively validated SenSet in human ex vivo lung tissue, using precision-cut lung slices (PCLS) subjected to senescence induction by bleomycin, doxorubicin, or irradiation^19–21^. This approach validated the superiority of SenSet in identifying SnCs and defined senescence cell-type gene signatures.

## Results

We developed a computational method that uses the largest publicly available single-cell lung dataset— the Human Lung Cell Atlas (HLCA)^18^—to identify senescent cell populations in the lung across different ages. Our approach is based on positive-unlabeled learning under covariate shift (PUc)^22^ that enables the derivation of a list of senescence markers via direct differential expression (DE) analysis of healthy (i.e., non senescent) and senescent cells (Fig. 1). To achieve this, we trained and tested this PUc learning approach by treating different age groups in the HLCA as (un)labeled data (Methods).

**Fig. 1:**
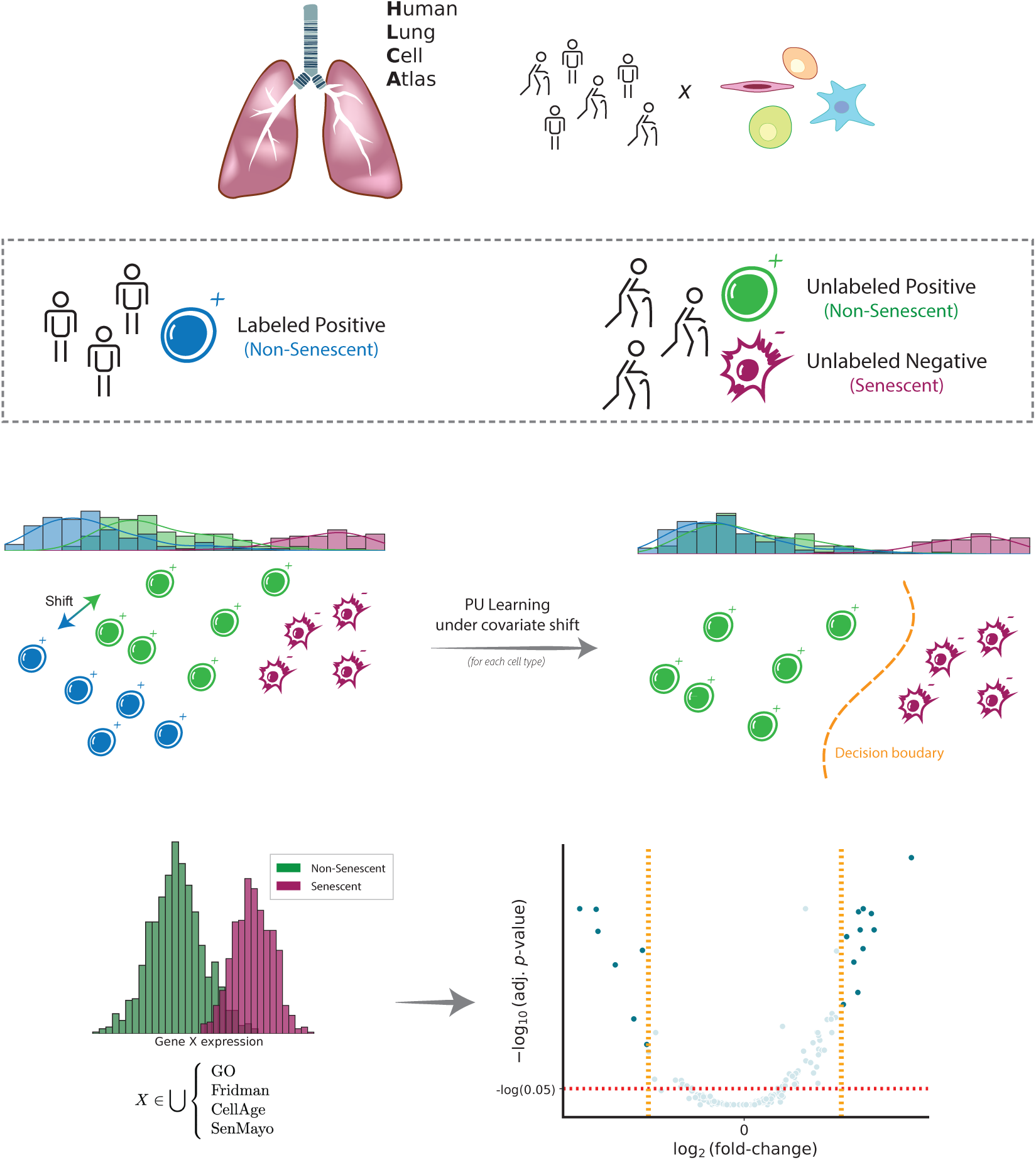
Schematic illustration of PUc learning for identifying SnCs. Given a large single-cell lung cohort from young and old individuals (top), we designate cells from young individuals as non-senescent cells (positive). Cells in older individuals are unlabeled initially. We then use positive-unlabeled learning under covariate shift (PUc) to identify SnCs in older individuals (middle). Using these cells, we develop an expression profile for senescence markers in several different cell types (bottom).

### Demographics and characterization of the HLCA

The 106 tissue samples in the HLCA were derived from individuals aged 10 to 76 years, including 51 never-smokers, 19 former smokers, and 28 active smokers (Fig. 2A). Overall, within the HLCA, tissue samples were derived from 69 males and 38 females. Among the 51 never-smokers, 33 were male with most of them belonging to the older age group (Fig. 2A,B). Thus, this dataset enables analysis of senescence signatures based on cigarette smoke exposure. Most of the samples originated from lung parenchyma from donor lungs that have been deemed not suitable for transplantation (Fig. 2C). A total of 50 cell types were present in the atlas at the finest annotation level (Fig. 2D-H), with respiratory basal cells and alveolar macrophages being the two most prevalent cell types among the never-smoker (NS) group (Fig. 2G). Among active smokers, type II pneumocytes and basal cells were the most common (Supp. Fig. 1). The total number of cells was 584,944 with 301,791 cells from never-smokers (Supp. Fig. 2). Total gene counts increased with age among smokers with a Pearson correlation of 0.33 (*p* = 0.01), while a slight but non-significant decrease was observed for never-smokers (Fig. 2I). We first analyzed the expression of *CDKN1A* and *CDKN2A*, which encode the senescence markers p21 and p16, respectively. Notably, *CDKN1A* was upregulated in smokers for the older two age groups (Fig. 2J). The most significant upregulation was observed for the oldest smoker group compared with non-smokers (two-tailed t-Test, *p* = 0.004). No significant differences in *CDKN2A* expression were observed across age or by comparing smokers with never smokers (Fig. 2J).

**Fig. 2:**
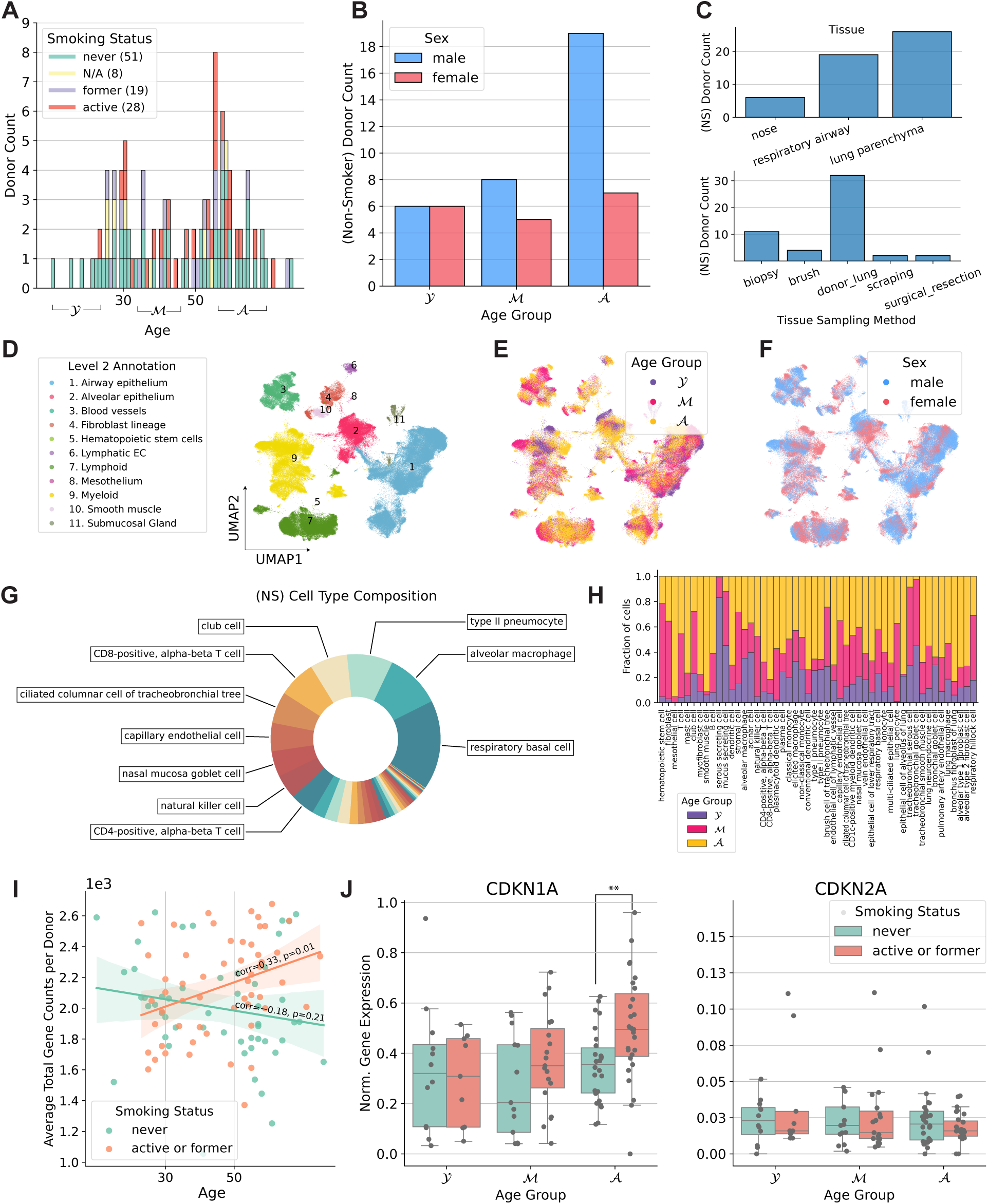
Summary of the Human Lung Cell Atlas. (A) Distribution of donors by age and smoking history. (B) Number of donors by age group and sex. (C) Number of donors broken down by tissue and sampling method (non-smokers). (D-F) UMAP plots describing level 2 cell types, age groups, and sex (NS = non-smoker). (G) Cell type proportion among non-smokers. (H) Age group representation across cell types (NS). (I) Average cell total counts for each donor. (J) Normalized gene expression values for *CDKN1A* and *CDKN2A*.

### Generation of SenSet from the HLCA

Several senescence gene sets have been published to date (GO:0090398, Fridman, SenMayo, and CellAge^12–14,17^). We first examined the extent of overlap between these gene sets. We observed that the pairwise overlap between them is relatively small compared with the size of each individual gene set, with the highest overlap of 34 genes shared between the GO and CellAge sets (Fig. 3B). The union **U** of all sets contains 501 unique marker genes, of which 434 were detected in the HLCA.

**Fig. 3:**
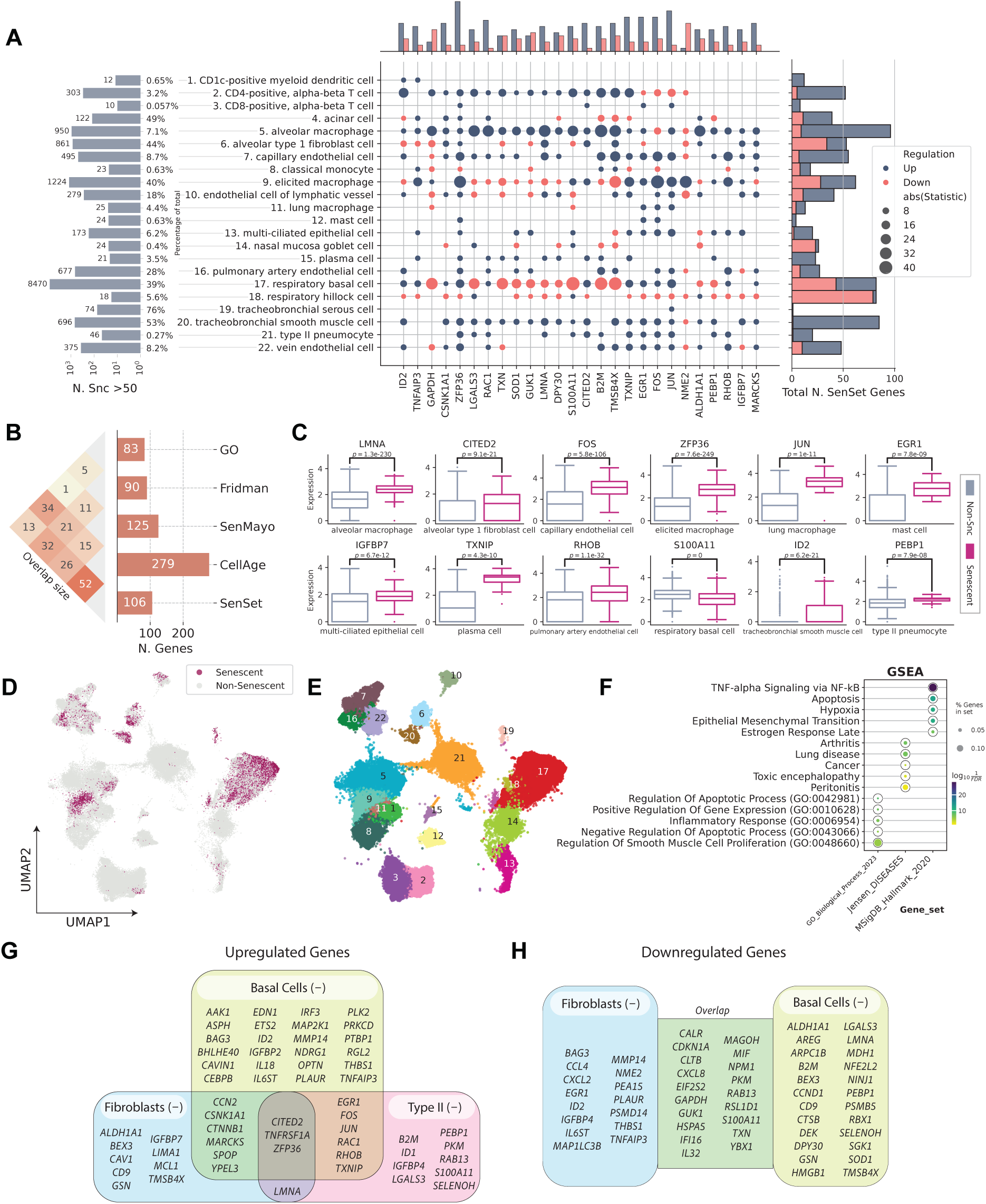
PUc identified a novel SenSet senescence signature. (A, left to right) The number and percentage of cells identified as senescent in A; the most frequent genes enriched for most cell types; the total number of SenSet marker genes assigned to a cell type. (for full figures see Supp. Fig. 3-7 or Supp. Tables) (B) Overlap sizes of SenSet with the prior lists. (C) Selected marker genes for some of the cell types and distribution among healthy and SnCs. (D) UMAP plot of senescent cells in A. (E) UMAP plot of cell types assigned at least one marker in A. The cell type numbers for each cluster correspond to the names listed in panel A. (F) Top GO, Jensen, and MSigDB terms enriched for SenSet. (G-H) SenSet genes enriched for basal^(−)^ cells, fibroblasts^(−)^, and type II pneumocytes^(−)^.

We sought to identify a subset of **U** that demonstrates greater sensitivity for senescence. The PUc estimator^22^ constructs a model of healthy cells based on data from (non-smoker) young and middle-aged individuals (groups Y, M, respectively) and applies this model to identify cells that are senescent in the aged group (group A, Fig. 1, 2A). PUc accounts for potential covariate shifts in A, which may arise due to other aging processes and hallmarks, including inflammation or epigenetic alterations^23,24^. The advantage of this approach is that it allows for a direct comparison of SnCs against non-SnCs within the same group—the oldest age group A. This addresses the challenge of age-related confounding factors that may arise when comparing older with younger individuals. While PUc identifies cells that generally deviate from the healthy (young) profile, we hypothesize that a significant proportion of these non-healthy cells identified by PUc are indeed senescent. We denote these cells with a (−) superscript to signify that they belong to the negative (senescent) class. Similarly a (+) superscript will denote non-senescent cells for that group.

We applied PUc to 31 cell types in the HLCA with a sufficient number of cells per age group (at least 50), using data exclusively from non-smokers to study senescence genes without the impact of cigarette smoke exposure. PUc identified at least 10 cells in the negative class—claimed here to be senescent—within 22 of these cell types (Fig. 3A,D,E). The proportion of cells assigned to this class ranged from 0 to 76 %. Major cell types with high enrichment for senescence markers included alveolar type 1 fibroblasts, respiratory basal cells, and tracheobronchial smooth muscle cells (44 %, 39 %, and 53 %, respectively). Of note, for some of the cell types with a high percentage of SnCs, such as tracheobronchial serous cells (76 %), few differentially expressed (DE) genes in these cell types were consistent with other types, suggesting mislabeling or that PUc assumptions for senescence might not hold true for this example.

### Differential expression analysis of SenSet genes

Next, we performed a DE test between the senescent 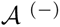 and non-senescent 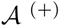 cells for each cell type within the old age group, and identified genes in the overlap senescence signature that were significantly enriched in at least six cell types (FDR = 0.05). This number was chosen to get a set of approximately 100 genes, which we termed SenSet (Table 1, Fig. 3G). SenSet showed the highest overlap with CellAge (52 out of 106 genes), followed by Fridman (32) and SenMayo (26) (Fig. 3B).

**Table 1:**
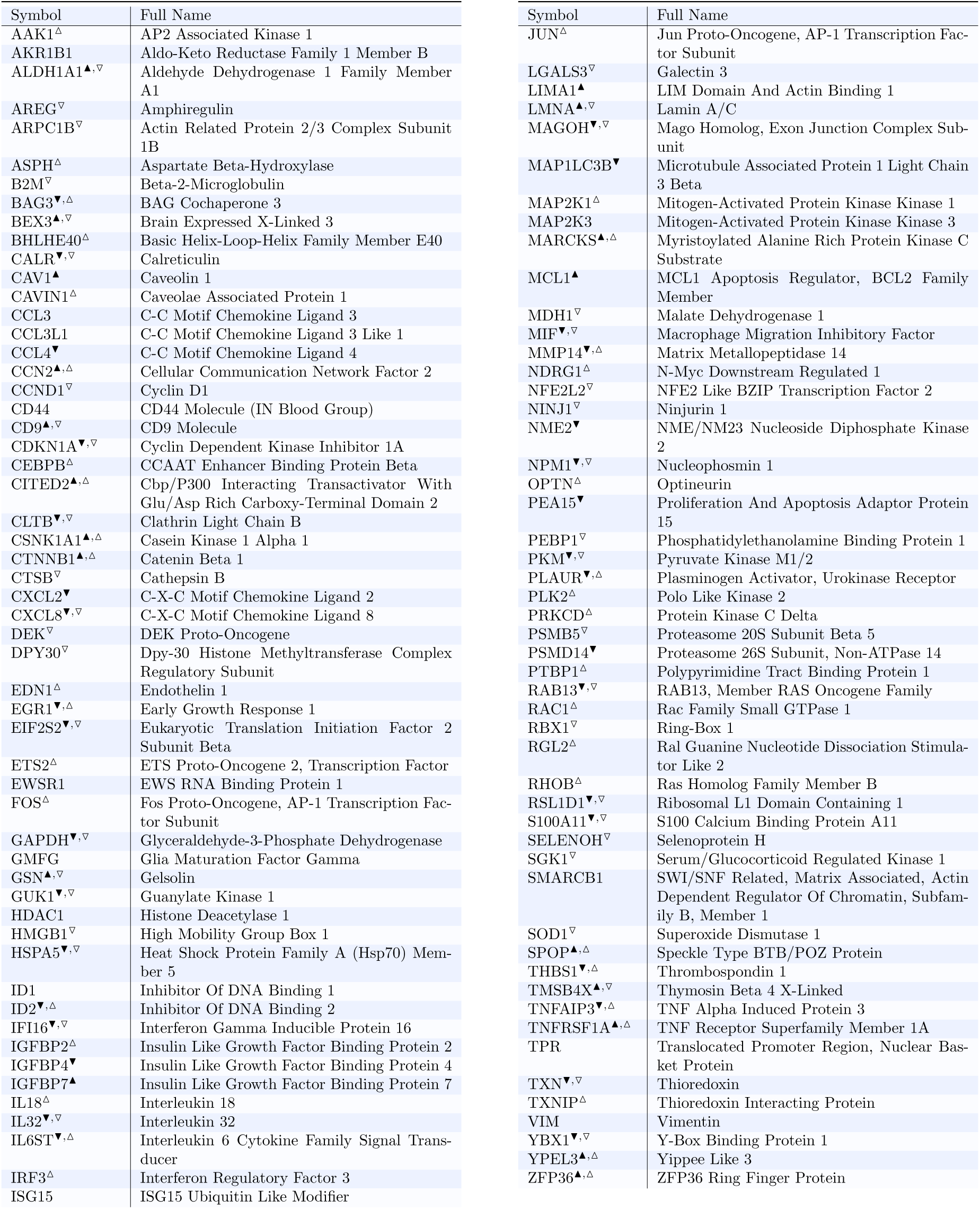
All 106 SenSet genes with full names. Marked are genes upregulated in fibroblasts^(−)^ (▲), downregulated in fibroblasts^(−)^, (▼), upregulated in basal^(−)^ cells (△), and downregulated in basal^(−)^ cells (▽) in the HLCA.

The senescence hallmark gene *CDKN1A* was enriched in 7 cell types, thus is included in SenSet, while *CDKN2A*, which was enriched in only alveolar macrophages, is not included. SenSet also contains SASP protein members, such as C-X-C motif chemokine ligand 8 (*CXCL8*), interleukin 18 (*IL18*), and insulin growth factor binding protein 7 (*IGFBP7*)^25–28^. Additional genes upregulated in most senescent cell types include ZFP36 ring finger protein (*ZFP36*, 16 cell types), Jun proto-oncogene (*JUN*, 13), and thioredoxin interacting protein (*TXNIP*), early growth response 1 (*EGR1*), Fos proto-oncogene (*FOS*) (11 each), all of which encode for proteins known to be involved in signaling in transcriptional response to hypoxia and cellular stress^29,30^. In contrast, nucleoside diphosphate kinase 2 (*NME2*), a suppressor of apoptosis, was downregulated in 9 cell types, followed by nucleophosmin 1 (*NPM1*) and glyceraldehyde-3-phosphate dehydrogenase (*GAPDH*) (7 each), involved in DNA replication and cell cycle^31–33^.

Gene set enrichment analysis (GSEA)^34^ using the MSigDB gene set^35^ revealed that SenSet is significantly enriched for genes involved in TNF-alpha signaling via NF-*κ*B (27 genes, *q* = 1e−29), apoptosis (15 genes, *q* = 1e−13), and hypoxia (15 genes, *q* = 1e−12) (Fig. 3F). Additionally, SenSet is enriched for genes associated with arthritis (12 genes, *q* = 1e−8) and lung disease (10 genes, *q* = 1e−8) based on Jensen’s disease set^36^. Notably, gene ontology (GO)^17^ analysis highlighted enrichment for the process “regulation of smooth muscle cell proliferation” (*q* = 1e−6).

### Cell type specific enrichment of SenSet genes

Not surprisingly, cell types displayed considerable heterogeneity in the expression of SenSet markers. From all SenSet genes, 87 were upregulated in alveolar macrophages^(−)^ and 84 in tracheobronchial smooth muscle cells (TSM)^(−)^, representing the highest numbers among the 22 cell types considered. The finding for TSM cells aligns with the GO analysis performed earlier. Conversely, basal^(−)^ and type 1 fibroblasts^(−)^ showed a downregulation of 43 and 34 genes, respectively (Fig. 3A,C, Table 1).

Type II pneumocytes and fibroblasts are crucial structural cell types in the lung that have been implicated in senescence^1,37,38^. In fibroblasts^(−)^, 19 SenSet genes were upregulated. Type II pneumocytes^(−)^ also show an upregulation of 19 SenSet genes (different set), and 1 downregulated gene, *CTNNB1*. We found substantial overlap in upregulated genes between basal^(−)^ cells and type II^(−)^ pneumocytes, fibroblasts^(−)^, respectively, with *TNFRSF1A*, *CITED2*, and *ZFP36* in common across all three. For instance, 9 genes were upregulated in both fibroblasts^(−)^ and basal^(−)^ cells, and 9 genes were also upregulated in both type II pneumocytes^(−)^ and basal^(−)^ cells (Fig. 3G).

Basal cells represent bona fide stem cells of the lung, and stem cell exhaustion—recognized as a hallmark of aging—has been associated with senescence^39^. Among 34 downregulated SenSet genes in fibroblasts^(−)^, 19 of these were also downregulated in basal^(−)^ cells (Fig. 3H).

Several genes were found to be upregulated in one cell type and downregulated in the other. For instance, *LMNA*, encoding for lamin A protein, known to be downregulated in more immature/undifferentiated cells^40^, was downregulated in basal^(−)^ cells, but upregulated in both fibroblasts^(−)^ and type II pneumocytes^(−)^.

### Senescence validation in human tissue model

To further refine and validate SenSet, we utilized a highly complex ex vivo human 3D tissue culture model based on precision cut-lung slices (PCLS). PCLS recapitulates the complexity of the lung environment in situ, enabling the study of various lung cell types and lineages present in the parenchymal region, along with the extracellular matrix (ECM) within the lungs’ native 3D architecture at high temporal and spatial resolution. We have previously demonstrated that this model can be applied to mimic the onset and progression of lung injury and diseases and enables testing potential therapeutics in living, diseased human tissue^41–43^.

PCLS were generated from the lower left lung lobe (Fig. 4A) of healthy donors aged 20 to 78 (Table 2). To induce cellular senescence, PCLS were treated with either bleomycin (15 µg*/*mL) or doxorubicin (0.1 µM) for up to six days. Lung structure remained intact after six days in culture, as shown by hematoxylin and eosin (H&E) staining (Figure 4B).

**Fig. 4:**
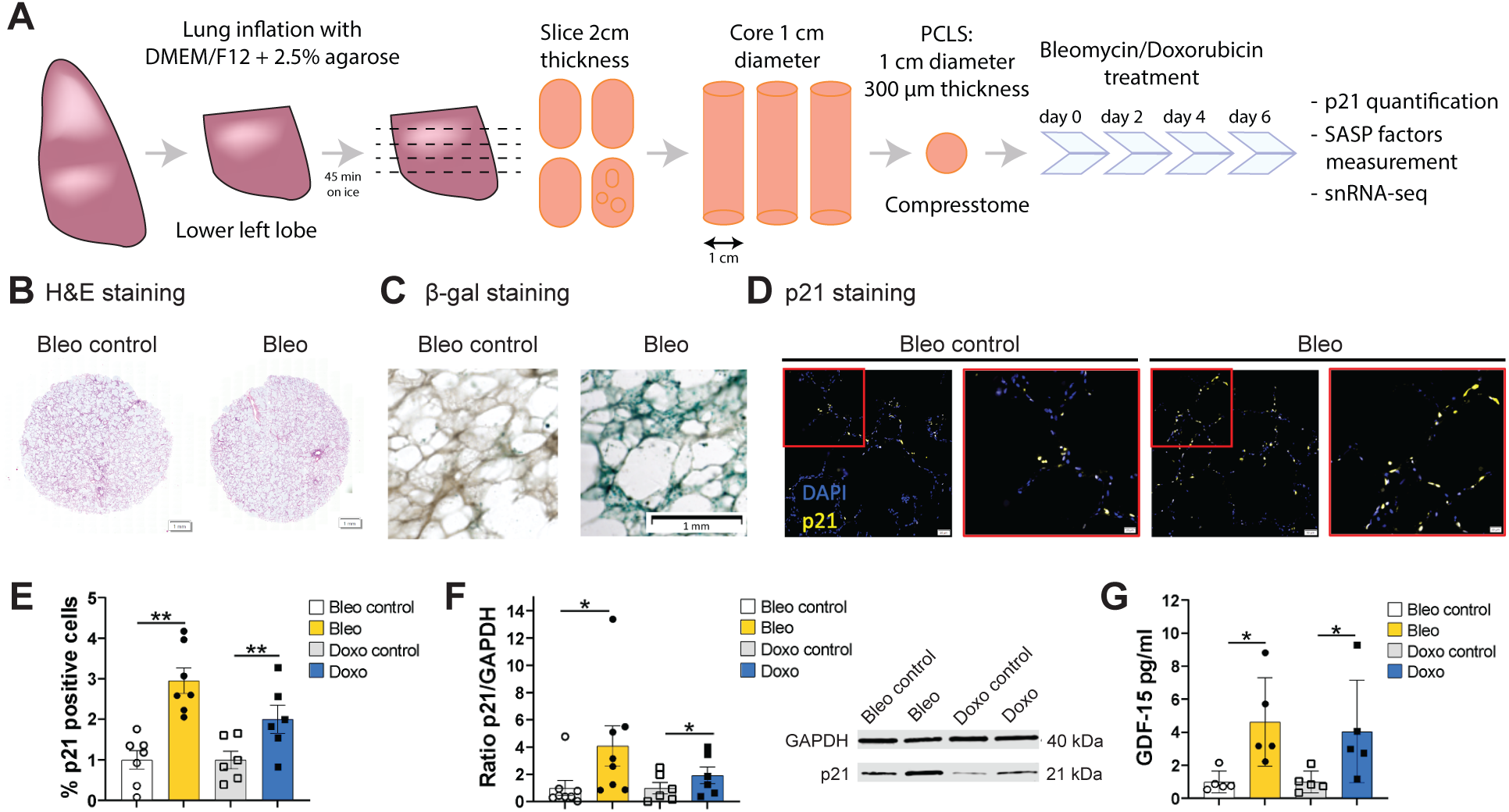
Senescence induction in human PCLS by DNA damage. (A) PCLS were generated from healthy donors lower left lobe lung with an age range of 20 to 78 years-old. Senescence was induced by treatment with bleomycin (Bleo) at 15 mg*/*mL, or doxorubicin (Doxo) at 0.1 µM for 6 days, and PCLS and supernatants were collected. (B) Hematoxylin eosin (H&E) staining on 4 µm sliced formalin-fixed paraffin embedded human PCLS at day 6. (C) *β*-galactosidase staining on whole PCLS at day 6. (D) p21 immunohistofluorescence (IHF) on 4 µm sliced formalin-fixed paraffin embedded human PCLS at day 6. (E) p21 positive cells quantification based on p21 IHF staining presented in (D) after bleomycin (*n* = 7) or doxorubicin (*n* = 6) treatment. (F) Quantification of p21 protein level by Western blot (WB) after bleomycin (*n* = 8) or doxorubicin (*n* = 6) treatment and representative blot. (G) SASP factor, *GDF-15*, measured by Luminex™ assay on human PCLS supernatants after bleomycin or doxorubicin treatment (*n* = 5). Paired t-test: \*\**p <* 0.005, \**p <* 0.05.

**Table 2:** Donor information. ^1^IHF: immunohistofluorescence; ^2^WB: western blot; ^3^Peritumor tissue; ^4^COPD Gold II; B: Bleomycin; D: Doxorubicin; I: Irradiation. If parentheses are missing, it is assumed to be (B, D).

| Donor | Sex | Age | Smoking status | IHF<sup>1</sup> p21 | WB<sup>2</sup> p21 | Multiplex assay | snRNA-seq |
| --- | --- | --- | --- | --- | --- | --- | --- |
| LTC 113 | F | 56 | Never | ✓ |  |  | ✓ |
| LTC 117 | M | 73 | Never | ✓ |  | ✓ | ✓ |
| LTC 118 | F | 23 | Never | ✓ |  | ✓(D) |  |
| LTC 119 | F | 38 | Never | ✓(D) |  | ✓(D) |  |
| LTC 120 | M | 21 | Never | ✓(B) |  | ✓(B) | ✓(B) |
| LTC 121 | M | 62 | Former | ✓(B) | ✓(B) | ✓(B) |  |
| LTC 124 | M | 36 | Never | ✓ | ✓ | ✓ | ✓ |
| LTC 127 | M | 41 | Never | ✓ | ✓ | ✓ |  |
| LTC 137 | F | 78 | Former | ✓ | ✓ |  |  |
| LTC 152 | F | 27 | N/A |  | ✓(B) |  |  |
| LTC 164 | M | 57 | Former |  | ✓ |  |  |
| LTC 176 | M | 25 | Never |  | ✓ |  |  |
| LTC 200 | M | 20 | Never |  | ✓ |  |  |
| E170<sup>3</sup> | M | 75 | Former |  |  |  | ✓(I) |
| E185<sup>3,4</sup> | F | 81 | Former |  |  |  | ✓(I) |
| E187<sup>3</sup> | M | 64 | Former |  |  |  | ✓(I) |
| E196<sup>3</sup> | F | 70 | Never |  |  |  | ✓(I) |

Senescence induction was validated by several readouts, with an increase in the number of *β*-galactosidase-positive cells (Fig. 4C) and p21-positive cells in PCLS (Fig. 4D, E). The percentage of p21 positive cells was significantly increased after bleomycin (2.9–fold ±0.8) and doxorubicin treatment (2.0–fold ±0.9, Fig. 4E). This was further confirmed by an increase in p21 after bleomycin (4.1–fold ±4.17) and doxorubicin treatment (1.9–fold ±1.49) (Fig. 4F). Moreover, *GDF-15*, a known SASP protein, was significantly increased after bleomycin (4.6–fold ±2.7) and doxorubicin treatment (4.1–fold ±3.1) (Fig. 4G).

To validate that our SenSet list—which was derived from the HCLA using lung tissue across the ages—is indeed a senescence signature, we subjected our human senescence induction model to single-nucleus RNA sequencing (snRNA-seq) and further analyzed cell type specific gene expression. Senescence was induced in PCLS by bleomycin or doxorubicin as described above and we further included PCLS subjected to irradiation, as previously reported^44^. The samples used for irradiation originated from peritumor tissue.

Importantly, we identified all major cell lineages in our ex vivo human tissue model and identified four major epithelial cell types (Fig. 5E) based on lung canonical markers^45^ (Supp. Fig. 8-9), with no discernible batch condition or cell cycle effects on data following integration using scVI^46^ (Fig. 5D).

**Fig. 5:**
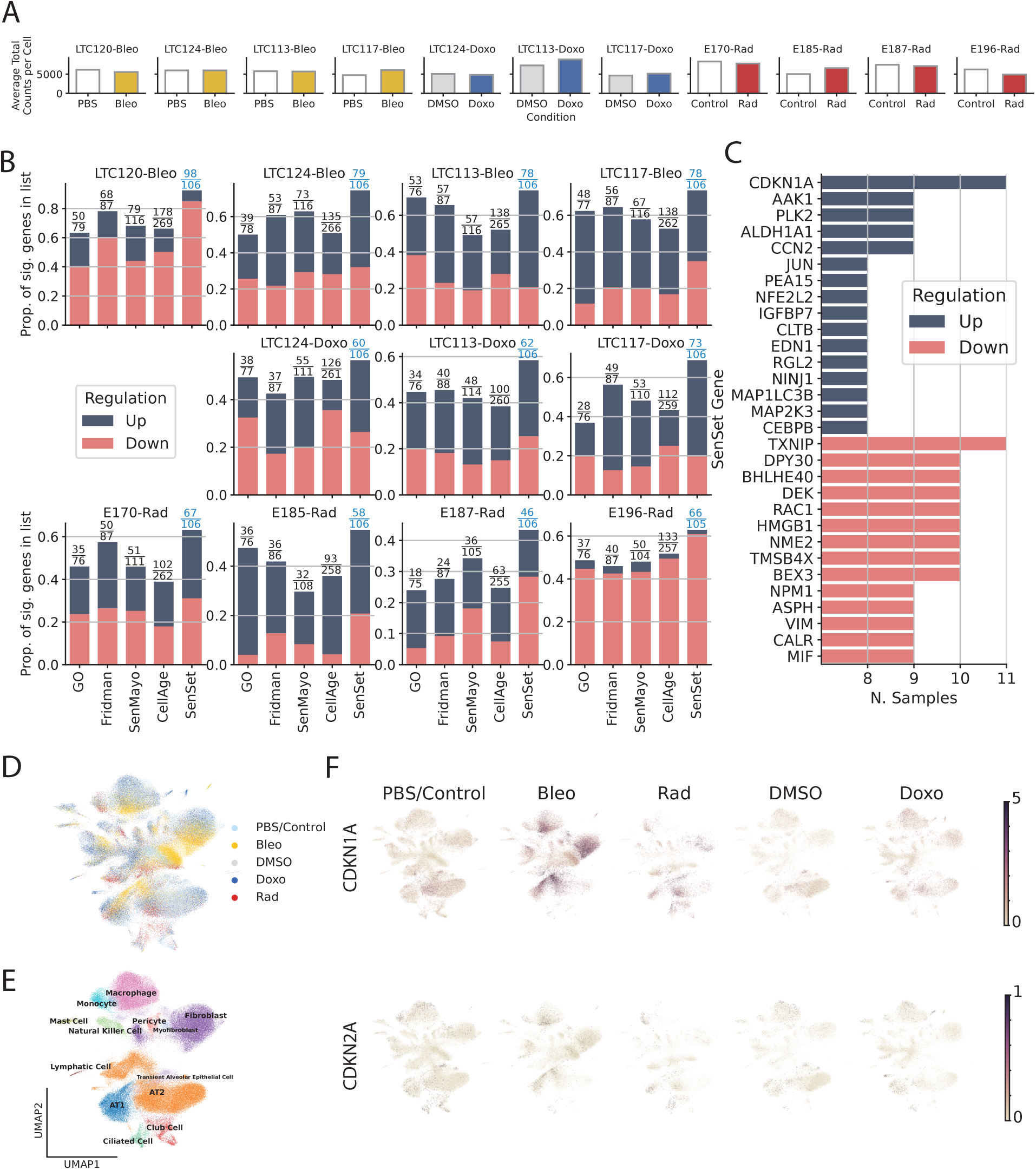
Validation of SenSet. (A) Average total counts per cell across samples and conditions. (B) For each subject, we show the fraction of the genes in each list which were significantly up or downregulated with treatment. (C) SenSet genes up (down)regulated in most samples. (D) UMAP plot of scVI integrated data. (E) Clusters identified using Leiden clustering on scVI embeddings. (F) Normalized expression of *CDKN1A* and *CDKN2A* across conditions.

For each gene in SenSet as well as each of the prior gene sets, we performed a rank sum DE test to determine if the gene is up or downregulated in our PCLS senescence models. Notably, SenSet achieves the highest proportion of significantly regulated genes in all samples when compared to all prior lung senescence lists (FDR = 0.05, Fig. 5B). Several SenSet genes that were upregulated in 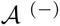 (HCLA) were also upregulated after senescence induction in human PCLS ex vivo, including *JUN*, and *IGFBP7*. The transcription factor *TXNIP*, which is known to be suppressed by p21 overexpression under disturbed shear stress in endothelial cells^47^, was downregulated in response to all three senescence inducers as well as the cell cycle regulators *NME2* and *NPM1*, which were also the top downregulated genes in 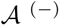 (Fig. 5C). *CDKN1A* was increased after all three treatments, with the highest induction after bleomycin treatment. No such increase was observed for *CDKN2A* (Figure 5C, F).

After we compared the expression of senescence markers across conditions using the entire sample, we next turned our attention to cell type-specific DE analysis of the marker genes, using the manually annotated cell types. For each cell type, we performed a similar rank sum DE test as before between treatment and control samples, and combined the *p*-values of these tests using Pearson’s method^48^ (Fig. 6A). This analysis revealed that SenSet markers showed significant enrichment across 10 cell types (*p* ≤ 0.05). In comparison, markers from Fridman and SenMayo lists showed significant enrichment in only four cell types. CellAge, on the other hand, was not significantly enriched in any cell type, likely due to the high number of non-DE markers in that list.

**Fig. 6:**
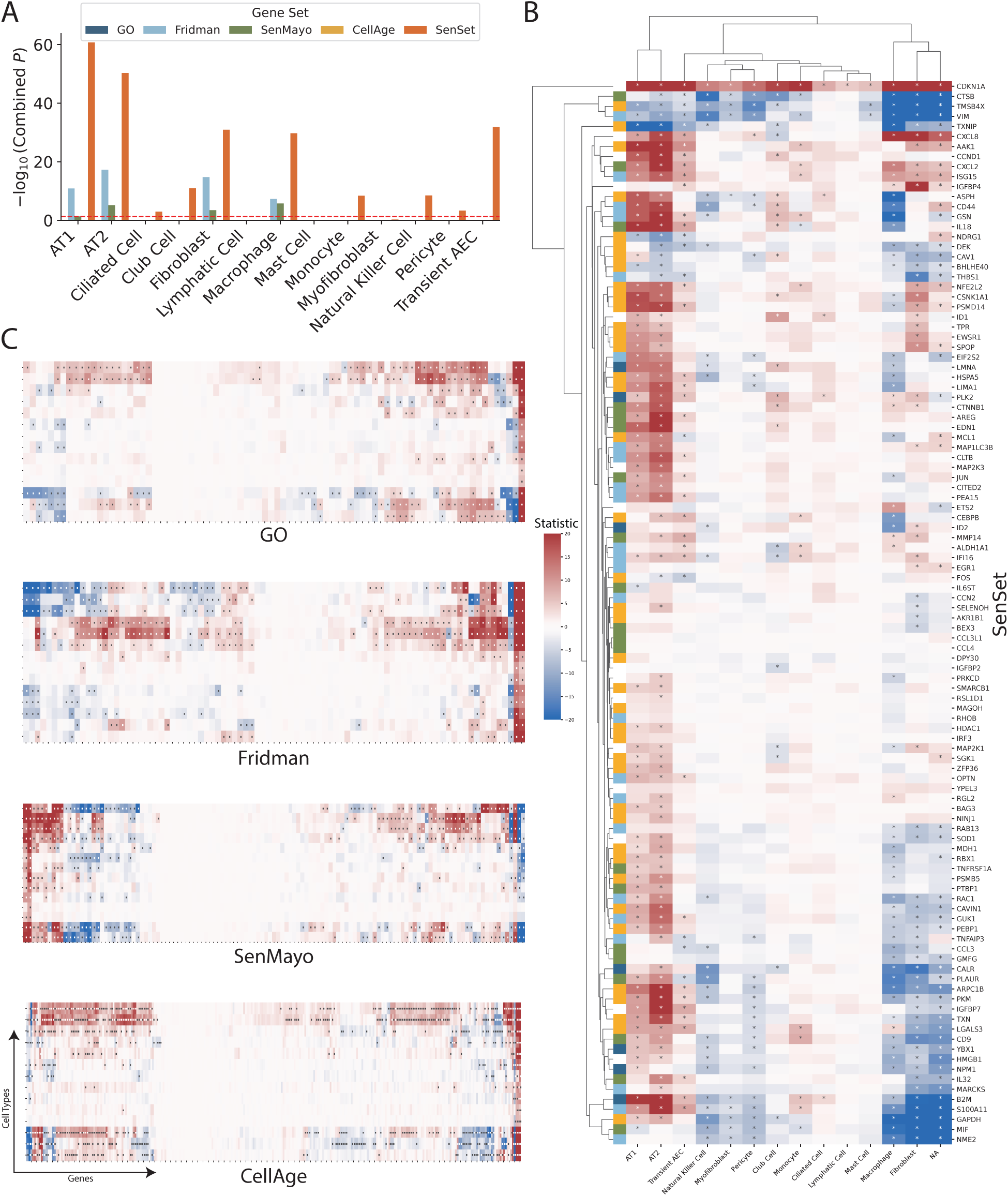
Cell Type-specific signatures. (A) For each cell type and gene set, we ran DE tests between the two conditions and show combined *p*-values (Pearson’s method) for each marker gene. (B-C) Rank sum test statistics for every marker gene and cell type in PCLS \**q* = 0.05 (for full figures see Supp. Fig. 10-13).

A close inspection of the markers across cell types confirmed that in the prior lists, many genes were not DE in any cell type (Fig. 6C). In contrast, nearly all SenSet genes (all but seven) were DE in at least one cell type (Fig. 6B). SenSet markers were predominantly upregulated in AT1^(−)^ and AT2^(−)^ cells, while mostly downregulated in fibroblasts^(−)^ and macrophages^(−)^. Notably, 15 genes downregulated in fibroblasts^(−)^ in the HLCA dataset were also downregulated in fibroblasts^(−)^ in the treated PCLS data: *CALR*, *GAPDH*, *GUK1*, *IL32*, *MIF*, *NME2*, *NPM1*, *PKM*, *PLAUR*, *RAB13*, *S100A11*, *THBS1*, *TNFAIP3*, *TXN*, *YBX1*. A similar correspondence was observed for (elicited) macrophages and AT2 cells. For AT2^(−)^ cells, 13 SenSet markers were upregulated in both the HLCA (19 total) and the PCLS (73 total), including *TNFRSF1A*, *PKM*, *PEBP1*, *ID1*, *ZFP36*, *LGALS3*, *RAC1*, *LMNA*, *S100A11*, *CITED2*, *B2M*, *JUN*, *SELENOH*.

### Analysis of smokers in the HLCA

Air pollutants and cigarette smoke exposure are a major risk factor for the development and exacerbation of age-related lung diseases, including chronic obstructive pulmonary disease (COPD). COPD incidence and prevalence increase with age^49–52^. The disease is characterized by impaired lung repair and progressive distal lung tissue destruction (emphysema) and airways remodeling and inflammation (chronic bronchitis)^53^. First, we computed the Wasserstein distance between pairs of smokers and non-smokers from different age groups *across all genes*. The Wasserstein distance is a measure of the difference between two probability distributions and provides a notion of how “close” the gene expression of two populations for a given cell type is. We found that for 12 out of 18 cell types, the expression space of young smokers (*<* 30) was closer in distribution to that of old non-smokers (≥ 50) than that of young non-smokers (Fig. 7A).

**Fig. 7:**
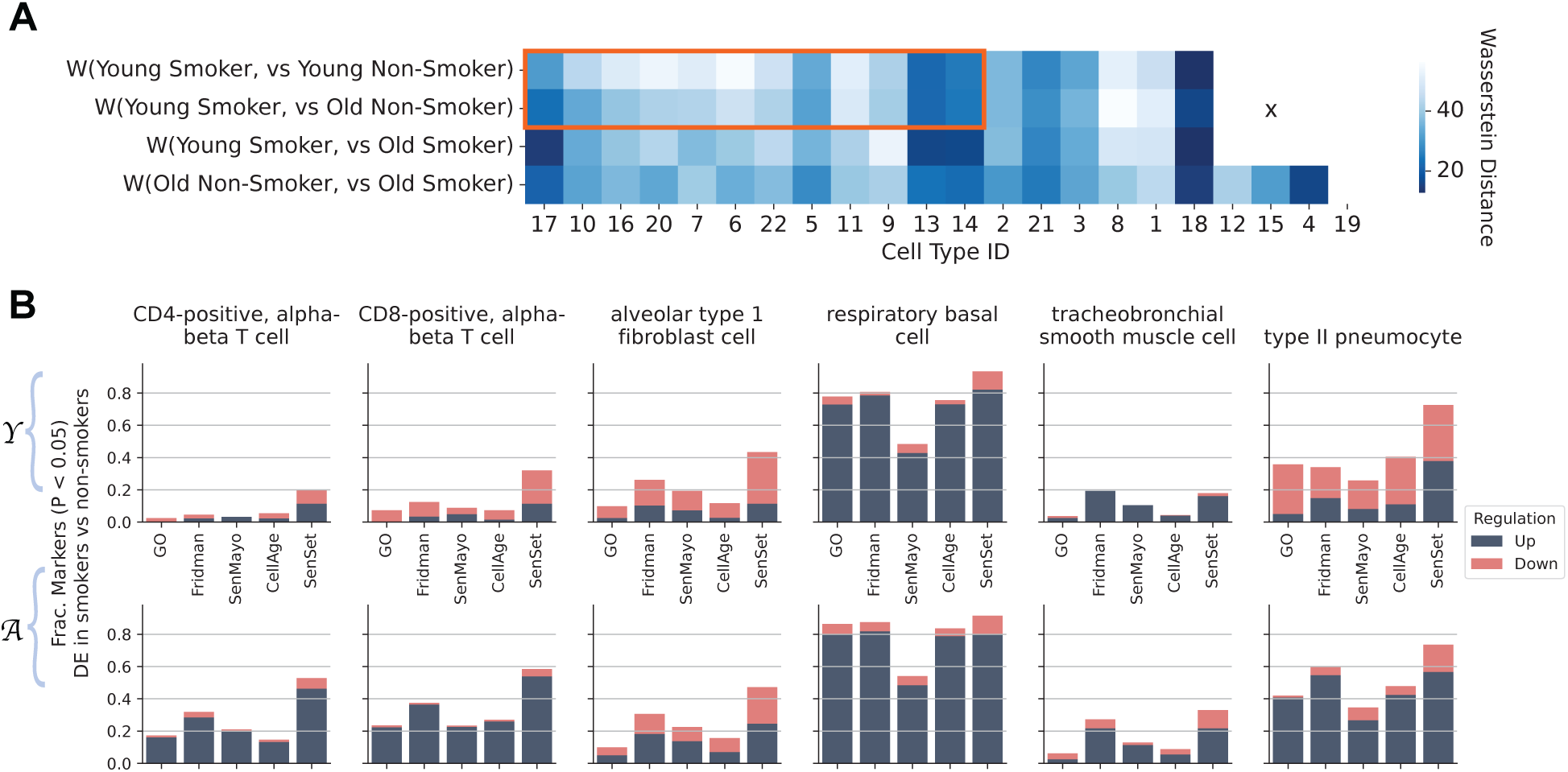
Comparisons of senescence marker genes between smokers and non-smokers in the HLCA. (A) Wasserstein distance between the gene expression profiles of smokers and non-smokers across different age groups. Cell types inside the red box exhibited a smaller distance for the pair (young smokers, old non-smokers) when compared to (young smokers, young non-smokers). (B) Fraction of genes enriched in smokers compared to non-smokers among young (Y) and old (A) patients for selected cell types.

Given that our SenSet signature was based on non-smokers, we next aimed to investigate whether SenSet enrichment is altered upon smoking. We performed DE testing on a few cell types of interest to evaluate if senescence markers were enriched in smokers for the youngest Y and oldest A age groups. The analysis showed differences in senescence enrichment between smokers of varying ages (Fig. 7B). Specifically, we found that most markers were upregulated in basal cells from smokers (around 80% of the genes across all lists except for SenMayo at 50%). For other cell types, including CD4/CD8-positive, alpha-beta T cells and AT2 cells, we found that aged smokers showed an upregulation of a higher number of senescence markers than young smokers, across all markers lists, with SenSet containing most such genes. Lastly, for AT2 cells around 35 % of all genes were downregulated and 40 % upregulated in the young, while around 60 % were upregulated in the older age group.

## Discussion

Aging remains the strongest risk factor for chronic lung diseases, with SnCs accumulating in tissues and contributing to pathology through mechanisms such as extracellular matrix remodeling and pro-inflammatory signaling. Cellular senescence is induced by diverse stimuli, including oncogene activation associated with tumor suppressor inactivation^54^, oxidizing agents inducing DNA damage^55^, or chemotherapeutic agents, such as bleomycin and doxorubicin^19,56^, with pathways varying by cell type, inducer, and time course. SnCs are defined by an altered DNA damage response (*γ*H2AX activation), expression of cyclin-dependent kinase inhibitors (p16, p21), enhanced SASP secretion (mTOR, cGAS–STING, NF–*κ*B), and apoptosis resistance (BCL-2)^57^, yet these features are not unique to senescence and often overlap with other cellular states including mitochondrial dysfunction or apoptosis.

This study presents a machine learning-based framework to identify a novel gene set specific to SnCs, originally generated using lung samples from the Human Lung Cell Atlas (HLCA). By leveraging single-cell transcriptomic data, our PU learning approach helped us distinguish SnCs across different age groups. Differential expression tests between SnCs and non-SnCs led to the creation of SenSet, which we validated in senescence-induced human lung ex vivo tissue. Senescence was induced in human PCLS by bleomycin, doxorubicin, or irradiation. We subjected our human senescence induction model to snRNA-seq to confirm the enrichment of SenSet at the cell-type level. SenSet is a subset of the union of four existing senescence marker sets, which enabled our weak supervision approach to identify senescence characteristics of cells with greater ease. Unlike prior lists, SenSet was entirely computationally-derived, which provides a data-driven, unbiased identification of genes that capture senescence-specific features across cell types.

Our findings revealed that fibroblasts and basal cells exhibited a high proportion of SnCs in aged lungs, accounting for 44 % and 39 % of all cells of that type in the oldest age group A. Fibroblasts contribute to alveolar maturation and regeneration by producing extracellular matrix components and are also central to the pathogenesis of age-related diseases, such as idiopathic pulmonary fibrosis (IPF), a disease which shows accumulation of senescent fibroblasts and increased SASP secretion^58^. Similarly, a recent work also demonstrated that fibroblasts are the main cell type undergoing senescence changes under homeostatic conditions and contribute to epithelial regeneration in a novel mouse model tracking p16^INK4a+^ cells^59^. Previous studies have shown that senolytic treatments targeting SnCs can alleviate fibrosis in mouse models^60^. Next to fibroblasts, basal cells showed significant SnC-associated gene regulation. Basal cells, which are the main stem cells in the proximal airways, can self-renew and differentiate into different airway epithelial cell types, such as secretory or ciliated cells^61^. These cells are critical for maintaining epithelial integrity and repair, particularly after injury. The number of basal cells gradually decreases in the proximal-distal axis in airway epithelium. Notably, stem cell exhaustion is a common hallmark of aging^62^.

Alveolar type 2 (AT2) cells, essential for surfactant production and alveolar repair, exhibited notable senescence-associated changes. AT2 cells have been demonstrated to harbor senescence features in the diseased lung such as in pulmonary fibrosis^37^. Elimination of senescent epithelial cells using senolytics has a beneficial effect in reducing lung fibrosis in mice^63^. Although only 0.27 % of AT2 cells in healthy HLCA samples were identified as senescent (Fig. 3), their transcriptional changes after senescence induction were among the most pronounced, with most SenSet genes upregulated after senescence induction in human PCLS. These results support the notion that AT2 cells are particularly sensitive to exposures and heavily involved in the injury response. In contrast, fibroblasts were the main cell type with most downregulated SenSet genes after senescence induction in PCLS (Fig. 6).

Several genes were consistently upregulated across fibroblasts^(−)^, basal^(−)^ cells, and AT2^(−)^ cells, emphasizing their shared senescence-associated pathways. Among the 19 upregulated genes in fibroblasts^(−)^, 8 genes are also upregulated in basal^(−)^ cells and/or in AT2^(−)^ cells such as *TNFRSF1A*, *YPEL3*, *SPOP*, *ZFP36*, *CITED2*, and *MARCKS* (Fig. 3G). Of these, *TNFRSF1A* encodes a receptor mediating inflammatory cytokine production, while *YPEL3*, a p53 downstream gene, is known to induce senescence^64^. *SPOP*, a tumor suppressor frequently mutated in cancers, was also upregulated and has been implicated in senescence induction and myofibroblast activation^65,66^. *ZFP36*, a gene regulating inflammatory cytokine production in psoriatic skin and metabolic pathways by growth factor induction,^67,68^, showed consistent expression patterns across SnCs (Fig. 3A), reflecting its broad involvement in cellular stress responses. *ZFP36* is known to be induced in cellular senescence in fibroblasts and in different human tissues^69^. Similarly, *CITED2*, which modulates TGF-*β* signaling^70^, was associated with cellular proliferation and senescence, with its reduced expression affecting aging and senescence in tendon-derived stem cells^71^. *MARCKS*, an actin-binding protein involved in cell motility and secretion, was also highly expressed in these cells.

A total of 19 genes were consistently downregulated in both fibroblasts^(−)^ and basal^(−)^ cells. For instance, *IL32*, a pro-inflammatory cytokine, was reduced in both, aligning with its role in regulating survival, proliferation and mitochondrial metabolism in myeloma cell lines^72^. *YBX1*, which regulates some SASP factors and senescence markers in human primary keratinocytes^73^, displayed similar downregulation in lung fibroblasts and basal cells.

In contrast, some genes were downregulated in fibroblasts^(−)^ while being upregulated in other cell types, highlighting cell type-specific senescence responses. For instance, *PLAUR*, involved in fibroblast-to-myofibroblast differentiation^74^, was downregulated in fibroblasts^(−)^ but upregulated in basal^(−)^ cells. Interestingly, *PLAUR* was identified as an upregulated gene in datasets from murine and human SnCs. *PLAUR* encodes for urokinase plasminogen activator receptor (uPAR), and treatment with uPAR-directed CAR T cells was a good senolytic strategy to decrease SnCs in vivo and in vitro^75^. Other genes included *ID2*, which is known to antagonize the growth-suppressive activities of p16 and p21^76^, and *TNFAIP3* which encodes for TNF-*α* induced protein 3 (A20). In mice, fibroblast A20 deletion recapitulates major pathological features of systemic sclerosis^77^. Altered expression of A20 in hematopoietic stem and progenitor cells leads to an aging-like phenotype, potentially impairing their functional capacity^78^. *IGFBP4* encodes for insulin-like growth factor binding protein 4, and is downregulated in fibroblasts^(−)^ and upregulated in AT2^(−)^ cells. IGFBP-4 was induced with irradiation in mice and humans and showed a pro-aging effect^79^.

*TXNIP*, associated with oxidative stress responses, was upregulated in both basal^(−)^ cells and AT2^(−)^ cells. *TXNIP* was shown to have a role in cellular senescence; its expression increases with age in *β*-cells and serum samples from humans, and it aggravates age-related and obesity-induced structural failure associated with an induction of cell cycle arrest and oxidative stress^80^. Furthermore, *TXNIP* deletion induces premature aging in hematopoietic stem cells inhibiting p38^81^. Other genes included *FOS* and *JUN* which are protooncogenes widely recognized for their involvement in the cellular senescence process^82,83^.

Validation of the SenSet gene list in an ex vivo model confirmed the induction of hallmark senescence markers, including p21. Among secreted proteins measured in the supernatants, GDF-15 exhibited the largest increase (Fig. 4G), aligning with its established role as an age-associated and senescence marker^84–86^.

Several SenSet genes were validated in our model, including *AAK1*, which induces several SASP factors^87^; *ALDH1A1*, linked to SASP regulation and senescence in ovarian cancer stem cells^88^; and *PLK2*, a kinase implicated in senescence pathways with reduced expression in glioblastoma^89^. *CNN2*, known to promote fibroblast senescence^90^, further supported the relevance of these markers in defining SnCs. Interestingly, *TXNIP*, which was upregulated in 11 cell types^(−)^ in HLCA, was downregulated in the ex vivo model, indicating its inducer-specific role in oxidative stress-mediated senescence. Similarly, genes such as *NME2* and *NPM1*, which play a role in cell cycle and tumor suppression^31,32^, were consistently downregulated in both datasets. This finding aligns with a prior study that *NPM1* upregulation inhibits p53-mediated senescence^91^.

Our findings underscore how environmental factors interact with aging processes to drive senescence in specific lung cell types. A comparative analysis of smoker and non-smoker groups within the HLCA revealed that smoking may accelerate aging in the lung. Notably, we shed light into major cell types that are likely mechanistically involved in these processes. Basal cells showed an upregulation of approximately 80 % of all SenSet genes, positioning them as primary responders to senescence in smokers.

Despite these results, our study has some limitations that need to be considered. The assumption that cells from individuals younger than 30 are universally healthy may not hold true in all cases, though the PUc classifier was robust against potential contamination by SnCs. Additionally, the use of arbitrary age thresholds (30 and 50) could influence the PUc classifier, although the framework remains valid as long as healthy cells in patients younger than 50 are similar. Recent studies have shown that significant developmental dysregulation occurs at the ages of 44 and 60^92^, leading us to believe that most healthy cells in this age range (30-50) are similar to those in the youngest group. The upper threshold was set to 50 in order to expand the set of individuals for this age group. Furthermore, the identification of marker genes is performed using one group only (ages 50+), and not by comparing age groups against each-other, so this likely does not constitute a major limitation of the subsequent analysis.

Some cell types, such as tracheobronchial serous cells exhibited an unusually large fraction of SnCs (76 %) as assigned by the PUc learner (Fig. 3A). However, only 1 marker was assigned to these cells, which may reflect mislabeling or insufficient data. Finally, only 18 of the 31 cell types analyzed using the PUc framework contained no SnCs in Y, suggesting that the method is robust to a small number of SnCs in the young.

In conclusion, this study advances our understanding of SnCs in the human healthy lung by identifying cell type-specific senescence markers applying a robust machine learning framework to a large aging cohort. The validation of these findings in ex vivo models with perturbed senescence strengthen their relevance, offering a foundation for future research into the role of SnCs in normal aging and accelerated aging in the context of chronic lung diseases, such as COPD. By linking cellular senescence to environmental stressors like smoking, this work highlights potential targets for therapeutic interventions aimed at mitigating the detrimental effects of senescence on lung health.

## Supporting information

SenSet.txt

## Acknowledgments

We gratefully acknowledge the provision of human biomaterial and metadata from the CPC-M bioArchive and its partners at the Asklepios Biobank Gauting, the LMU Hospital and the Ludwig-Maximillians University Munich. In addition, we thank the Center for Organ Recovery & Education (CORE). We are grateful to all Donors and their families for their support.

This research was supported by grants from the National Institutes of Health, including NIHU54AG075931 and U24CA268108.

## Data Availability

Data is available upon reasonable request.

## Declaration of Interests

The authors declare no conflict of interest.

## Code Availability

The code used for experiments in this paper is available at https://github.com/euxhenh/SenSet. The code for PUc learning is included with permission from the authors.

## Methods

### The Human Lung Cell Atlas

The (core) Human Lung Cell Atlas (HLCA)^18^ was downloaded from https://data.humancellatlas.org/hca-bio-networks/lung/atlases/lung-v1-0. Counts were already normalized. The HLCA harmonizes scRNA-seq data from 14 datasets, encompassing 106 individuals aged between 10 and 76 years. We removed one individual for whom the age was not available. While five levels of annotation are available in the data, we used the finest level assigning one of 50 cell types to over 500,000 cells for the analysis in this study. The dataset also contains individuals with a smoking history, including 19 former and 28 active smokers. Smoking status is not available for 8 individuals and these were not included in the analysis (Fig. 2).

### PCLS culture and senescence induction (Bleomycin and Doxorubicin)

Lungs from healthy donors (ages 20-78) were collected, and the lower left lobes were inflated with 2.5 % agarose in DMEM/F-12 with HEPES (Thermo Fisher Scientific Cat#12400024). After 45 minutes on ice, the lobes were sliced, and 1 cm diameter cores were extracted. Precision-cut lung slices (PCLS) were prepared using a Compresstome® to obtain 300 µm thick tissue slices with a diameter of 1 cm. The PCLS were then placed in a 24-well plate containing 1 mL of DMEM/F-12 medium supplemented with 1 % FBS, 1 % Penicillin-Streptomycin (Merck MilliporeSigma, Sigma-Aldrich Cat#P00781), and 0.3 µg*/*mL Amphotericin B solution (Merck MilliporeSigma, Sigma-Aldrich Cat#A2942) and incubated for 24 hours at 37 °C with 5 % CO2 (day −1).

At 24 hours (day 0), 72 hours (day 2) and 120 hours (day 4), the medium was replaced with fresh medium containing treatments diluted in DMEM/F-12 with 0.1 % FBS, 1 % Penicillin-Streptomycin, and 0.3 µg*/*mL Amphotericin B. The PCLS were treated under the following conditions: in PBS as the control, with 15 µg*/*mL bleomycin (Fresenius Kabi Cat#10361), with DMSO diluted at 1:100,000 in medium (Merck Millipore Sigme, Sigma-Aldrich, Cat#D2438), or with 0.1 µM doxorubicin hydrochloride (Merck Millipore Sigma, Sigma-Aldrich, Cat#D1515-10MG), originally dissolved in DMSO at 10 mM. After 168 hours (day 6), the supernatants were collected and frozen at −80 °C for future multiplex immunoassay analysis by Luminex™. The PCLS were snap-frozen in liquid nitrogen for future protein extraction and snRNA-seq using the 10x Genomics™ platform. Additionally, PCLS were fixed for 30 minutes in 4 % formaldehyde diluted in PBS (Life Technologies Cat#28908), washed in PBS, and embedded in paraffin. Separate PCLS were fixed for 30 minutes in the fixative solution from the *β*-galactosidase staining kit (Cell Signaling Technology Cat#9860) and then washed in PBS.

### PCLS culture and senescence induction (Irradiation)

Peritumor control tissue from three non-chronic lung diseases (N-CLD) patients and one COPD patient were obtained from the CPC-M bioArchive at the Comprehensive Pneumology Center (CPC Munich, Germany). Patients were two males and two females and had a mean age of 72.5 years old. Human lung tissue was filled with 3 % of low gelling temperature agarose in DMEM/F-12 (Thermo Scientific, USA) with phenol red supplemented with 0.1 % FCS, 1 % P/S and 1 % amphotericin B and kept at 4 °C for at least 1 hour. 500 µm PCLS were generated using either a vibratome HyraxV50 (Zeiss, Germany) or 7000smz-2 Vibratome (Campden Instruments, England). The day after slicing, fresh medium was added and PCLS were exposed to ionizing radiation using the RS225 X-ray cabinet (Xstrahl, Camberley, UK). Dose was calculated according to exposure time (30 Gray (Gy) = 12 min 24 sec) at 195kV and 15mA. Then, PCLS were kept in culture for up to 5 days at 37 °C, 5 % CO2 and medium was changed every 2-3 days.

### Ethic Statement

The study was approved by the local ethics committee of the Ludwig-Maximilians University of Munich, Germany (Ethic vote 19-630). Written informed consent was obtained for all study participants.

### Histology and immunohistostaining

Paraffin-embedded PCLS were sliced at a thickness of 4 µm. One slide was subjected to H&E staining, and another slide was used for p21 immunostaining. The slides were rehydrated through a series of baths in xylene, followed by 100 %, 95 %, 85 % ethanol, and finally water. The slides were then incubated at 105 °C for 20 minutes in 1X DAKO high pH buffer (Agilent Technologies, Dako Target Retrieval Solution pH 9 10X, Cat#S236784-2). Then the slides were washed in buffer A from Duolink® In Situ Wash Buffers (MilliporeSigma, Sigma-Aldrich Cat#DUO82049-20L), followed by incubation in 300 mM glycine for 30 minutes, and then in PBS containing 0.1 % Tween and 0.5 % Triton for 15 minutes.

The slides were then incubated with a blocking solution consisting of 2 % BSA, 0.1 % Triton, and 0.1 % Tween at 37 °C for 45 minutes, followed by an overnight incubation at 4 °C with the primary antibody, anti-p21 antibody [EPR362] (Abcam, Cat#ab109520), diluted 1:500 in the blocking solution. After washing in buffer A, the slides were incubated for 1 hour at room temperature with a 1:1000 dilution of the secondary antibody, anti-rabbit IgG 647 (Biotium, Cat#20047). Following further washes in buffer B from Duolink® In Situ Wash Buffers, the nuclei were stained with DAPI. The slides were then washed in buffer B and mounted with Fluoroshield Mounting Medium (Abcam, Cat#ab104135). Images were captured using an IX83 Olympus microscope, acquiring the entire PCLS area at 20x magnification.

Quantification of p21-positive cells was performed using Fiji macros, where the total cell number was determined by DAPI staining, and the number of p21-positive cells was identified by the nuclear p21 signal overlapping with DAPI staining. The percentage of p21-positive cells was calculated as the ratio of p21-positive cells to the total number of cells in each image.

### Protein extraction and western blotting

Four PCLS were sonicated three times at 35 % amplitude for 10 seconds in 200 µL of buffer (TPER buffer, Thermo Scientific, Cat#78510, supplemented with Halt™ Protease and Phosphatase Inhibitor Cocktail, EDTA-free (100X), Thermo Scientific, Cat#1861281, to a final concentration of 1X) and kept on ice. The samples were then homogenized for 15 seconds using a Thermo Fisher Scientific homogenizer and centrifuged at 300×g for 5 minutes at 4 °C. The supernatants were transferred to new 1.5 mL tubes and centrifuged at 10,000 rpm for 10 minutes at 4 °C. The supernatants were gently collected, and protein concentrations were quantified in triplicate using the Pierce detergent Compatible Bradford Assay kit (Thermo Scientific, Cat#23246), with 150 µL of reagent and 5 µL of sample per well. A standard curve was generated using the Prediluted Protein Assay Standards BSA Set (Thermo Scientific, Cat#23208). Absorbance was measured at 595 nm using SpectraMax ABS Plus Molecular Devices with SoftMaxPro 7.2.

Protein extracts were denatured in 1X Protein Loading Buffer (Li-COR, Cat#928-40004) containing 0.1 M dithiothreitol (DTT) (Sigma-Aldrich, Cat#43816) for 10 minutes at 95 °C. A total of 10 µg of denatured protein was loaded onto Criterion TGX long shelf-life Precast Gels (4-15 %, Bio-Rad, Cat#5671083) using 1X Tris/Glycine/SDS buffer (Bio-rad, Cat#1610772). The proteins were then transferred onto an Odyssey Nitrocellulose Membrane (Li-COR, Cat#926-31092).

The membrane was then blocked for 1 hour at room temperature in a 1:1 mixture of 1X TBS and Intercept Blocking Buffer (Li-COR, Cat#927-70001). The blocked membrane was incubated overnight at 4 °C with p21 antibody (Abcam, Cat#ab109520) diluted 1:1000 or GAPDH antibody (Abcam, Cat#ab9485) diluted 1:2500 in a 1:1 mixture of 1X TBST and Intercept Blocking Buffer.

The membrane was then washed three times for 10 minutes each with TBST and incubated with secondary antibodies: anti-rabbit red (IRDye® 680RD Donkey anti-Rabbit IgG, Li-COR, Cat#926-68073), diluted 1:20000 in a 1:1 mixture of 1X TBST and Intercept Blocking Buffer for 1 hour at room temperature. Images were captured and analyzed using an Odyssey imaging system with Image Studio software.

### SASP assessment by multiplex immunoassay

PCLS supernatants were harvested after six days in culture and stored at −80 °C. The multiplex immunoassay was performed using the Luminex™ platform, following the manufacturer’s instructions (Bio-Techne).

### Single-nucleus RNA sequencing by 10x Genomics™

PCLS were snap-frozen in cryotubes using liquid nitrogen after six days of culture and stored in liquid nitrogen. Four PCLS per experimental condition were then used for snRNA-seq, following the manufacturer’s instructions (10x Genomics™). Some minor individual differences in total-counts-per-cell were found across conditions (Fig. 5A).

### Identifying senescent cells in aged individuals via PU learning under covariate shift

Positive-Unlabeled (PU) learning is a variant of binary classification where the goal is to distinguish between positive and negative samples, with the restriction that only positive samples are seen during the training phase^16^. More concretely, the real data distribution is a mixture given by

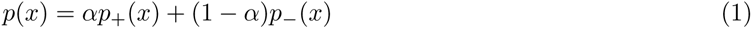

where *α* is the mixture proportion, and *p*_+_, *p*_−_ are the probability density functions of the positive and negative samples, respectively. In traditional binary classification, we are given training data 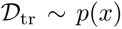 and test data 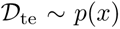. However, in PU learning, 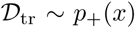 which makes it more challenging. Many methods have been proposed which typically assume some form of smoothness or separability of the classes, or treat the negative samples as noise^93–95^.

PU learning fits our setup as follows. Given the current uncertainty in identifying senescent cells (SnCs) using gene expression data, we treat the population of SnCs as negative samples within an unlabeled set. The objective is to recover a PU classifier capable of distinguishing SnCs from healthy cells. Subsequently, differential expression (DE) analysis between these two groups can be used to identify marker genes.

To obtain (labeled) training and (unlabeled) test sets for PU learning, we rely on different age groups present in the data, where 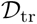 models young individuals, and 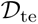 models more senior ones. To accomodate the PU learning setup, we require the following assumptions to hold: a) the young group contains few to no SnCs, b) the negative (non-healthy) class in the senior group consists mostly of SnCs, and c) the healthy cells in the senior group come from the same distribution as the healthy cells in the young. The last assumption is problematic as literature has shown that aging patients suffer from other hallmarks of aging not related to senescence such as inflammation, epigenetic alterations, or mitochondrial dysfunction^23,24,96^. Stated differently:

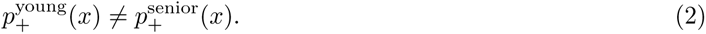

Therefore, we face a covariate shift^97^. In covariate shift, the dependence of the response variable *y* (i.e., senescence status) on gene expression *x* is the same for both the training and test sets, however, the input distributions may not be:

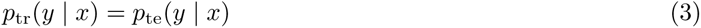

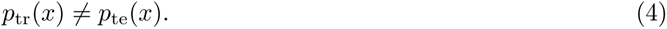

To address this, the PU learner needs to accommodate a covariate shift. Here, we rely on the formulation of Sakai & Shimizu which propose an unbiased risk estimator for covariate shift adaptation on PU learning, termed PUc^22^.

PUc assumes we are given three sets of samples: labeled training data 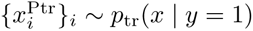, unlabeled training data 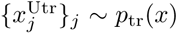, and unlabeled test data 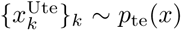. When covariate shift occurs, the PU risk on the test distribution *p*_te_ differs from the PU risk on the train distribution *p*_tr_. PUc addresses this by importance-weighting, where the ratio between test and train densities *w*(*x*) := *p*_te_(*x*)*/p*_tr_(*x*) is used to weight each sample during the computation of the risk^98,99^. In this case, the PUc risk becomes

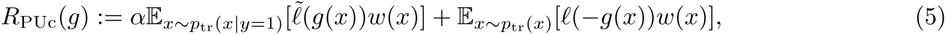

where *g* is a classifier and *ℓ* is a loss function with 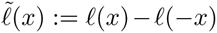. The PUc risk on training data can be shown to be an unbiased estimator of the PU risk on test data. The mixture proportion *α* is estimated from prior knowledge. Similar to the original work, we employ a linear-in-parameter classifier with a Gaussian kernel basis function. For more details on PUc, please refer to Sakai & Shimizu^22^.

### Deriving SenSet from the Human Lung Cell Atlas

During the gene set generation step, we kept only individuals without a smoking history from the HLCA (approximately 300,000 cells), to minimize potential confounding effects on the results. As described in the previous section, the PUc estimator requires three sets of samples: positive training samples, and unlabeled training and test samples. We estimated positive samples from individuals under the age of 30, assuming that the prevalence of SnCs in this group is minimal. The unlabeled training samples were estimated from individuals aged 30 to 50. Finally, the test samples were obtained from older individuals aged 50 and above, which is when covariate shift occurs. We call these three age groups 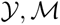, and 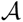, respectively (Fig. 2A, E). Cell types with fewer than 50 cells in any age group were excluded from the analysis.

To prepare the data for PUc learning, we first applied principal component analysis (PCA) independently for each cell type. We restricted the gene set used for PCA to the union **U** of all four existing senescence gene sets. By incorporating all known senescence-associated genes, we aim to achieve a “weak” separation of healthy cells and SnCs, which can be leveraged by the PUc learner. The top 10 components were used as training data for the PUc classifier. For all experiments, we set the mixture proportion *α* to 0.9, based on the prior assumption that approximately 10% of the cells are senescent. However, the estimator was robust to this value and returned percentages in the range 0 − 40%.

Given the inherent randomness of the algorithms used, some variability across runs is expected. Nonetheless, we observed that approximately 80-90 % of SenSet genes were consistently selected across runs with different random seeds.

Differential expression analysis is performed exclusively on the oldest age group, comparing healthy cells with SnCs. This approach is in contrast with methods that compare old and young individuals, where other aging signatures could introduce confounding factors. By directly comparing these two cell populations within the oldest age group, the analysis is specifically focused on senescence. A two-sided Wilcoxon rank-sum test was used to determine differentially expressed (DE) genes (FDR= 0.05). We tested only genes that belong to **U**. FDR-adjusted *p*-values were obtained using the Benjamini & Hochberg procedure^100^. Cell types with fewer than 20 SnCs were not considered due to the limited sample size. We selected DE genes that were enriched in at least six cell types (either up or downregulated), resulting in a set of 106 genes that constitute SenSet. A detailed table of all cell-specific SenSet genes and their test statistics is provided in the supplementary material.

GSEA^34^ was performed using the GSEApy package^101^. To compute the Wasserstein distances in Fig. 7A, we sampled at random 5000 cells for cell types with too many cells to speed up computation. Scanpy was used in some parts of the analysis including clustering and dimensionality reduction^102^.

### Integrating PCLS data and validating SenSet

For the overall sample comparison presented in Fig. 5B-C, we performed basic cell and gene filtering for all 11 samples. Cells with fewer than 500 total counts and fewer than 400 expressed genes were excluded. Genes with fewer than 50 total counts were also removed. Next, we normalized the total counts of cells. Normalized gene counts between treatment and control samples were compared using a two-sided Wilcoxon rank-sum test (FDR= 0.05) for each gene set.

For the cell-specific analysis, we first integrated the data using scVI^46^ focusing on the top 2000 variable genes. We used 2 hidden layers with 1000 nodes each. The dimensionality of the latent space was set to 30. Since we used raw counts for scVI, the gene likelihood was modeled as a zero-inflated negative binomial distribution. Nearest neighbors were computed in the scVI latent space, and clusters were identified via Leiden clustering^103^. Clusters were manually annotated based on canonical markers of lung cell types^45^. Clusters where no marker was significantly expressed were excluded from the cell-specific analysis. Wilcoxon rank-sum tests were used to assess whether SenSet genes were enriched in treatment samples compared to controls. For this analysis, bleomycin, doxorubicin, and irradiation cells were combined into one group.

For each gene set, we obtain a list of *p*-values for each gene based on the DE test. In order to perform a meta-analysis, we combine these *p*-values for each set using Pearson’s method, which emphasizes larger *p*-values. I.e., given a set of *p*-values 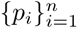, Pearson’s method computes the statistic

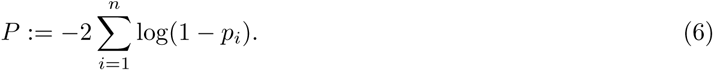

Under the null hypothesis *H*_0_ : *p_i_*~ *U* [0, 1], *i* = [*n*], the test statistic *P* follows a *χ*^2^ distribution with 2*n* degrees of freedom^48^. The combined *p*-value provides an overall assessment of the enrichment of a gene set within a given cell type.

## Notes

### Competing Interest Statement

The authors have declared no competing interest.

### Summary of Updates

Fixed author name; Added funding information; Added the full gene set as a text file.

