## Supplement.pdf for "SenSet, a novel human lung senescence cell gene signature, identifies cell-specific senescence mechanisms"

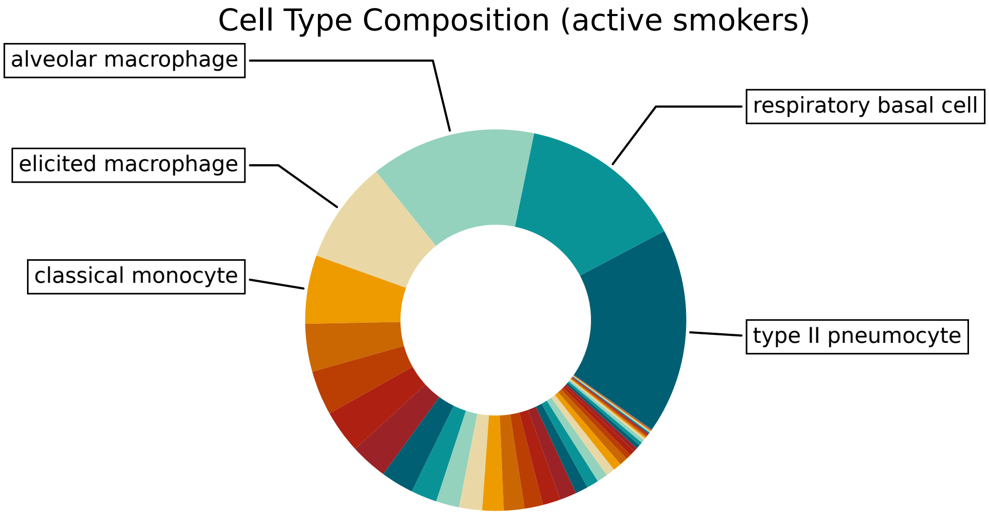


Figure 1: **Cell type proportion among active smokers.**

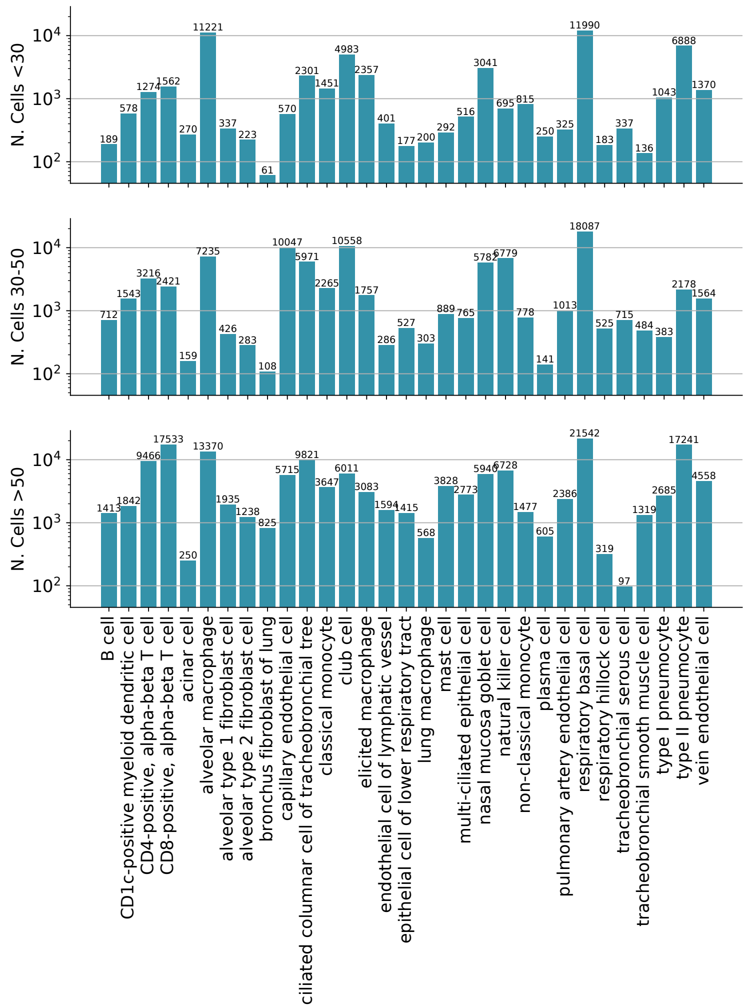


Figure 2: **Number of cells for each cell type used for PU learning and each age group in the HLCA dataset. Data from non-smokers only.**

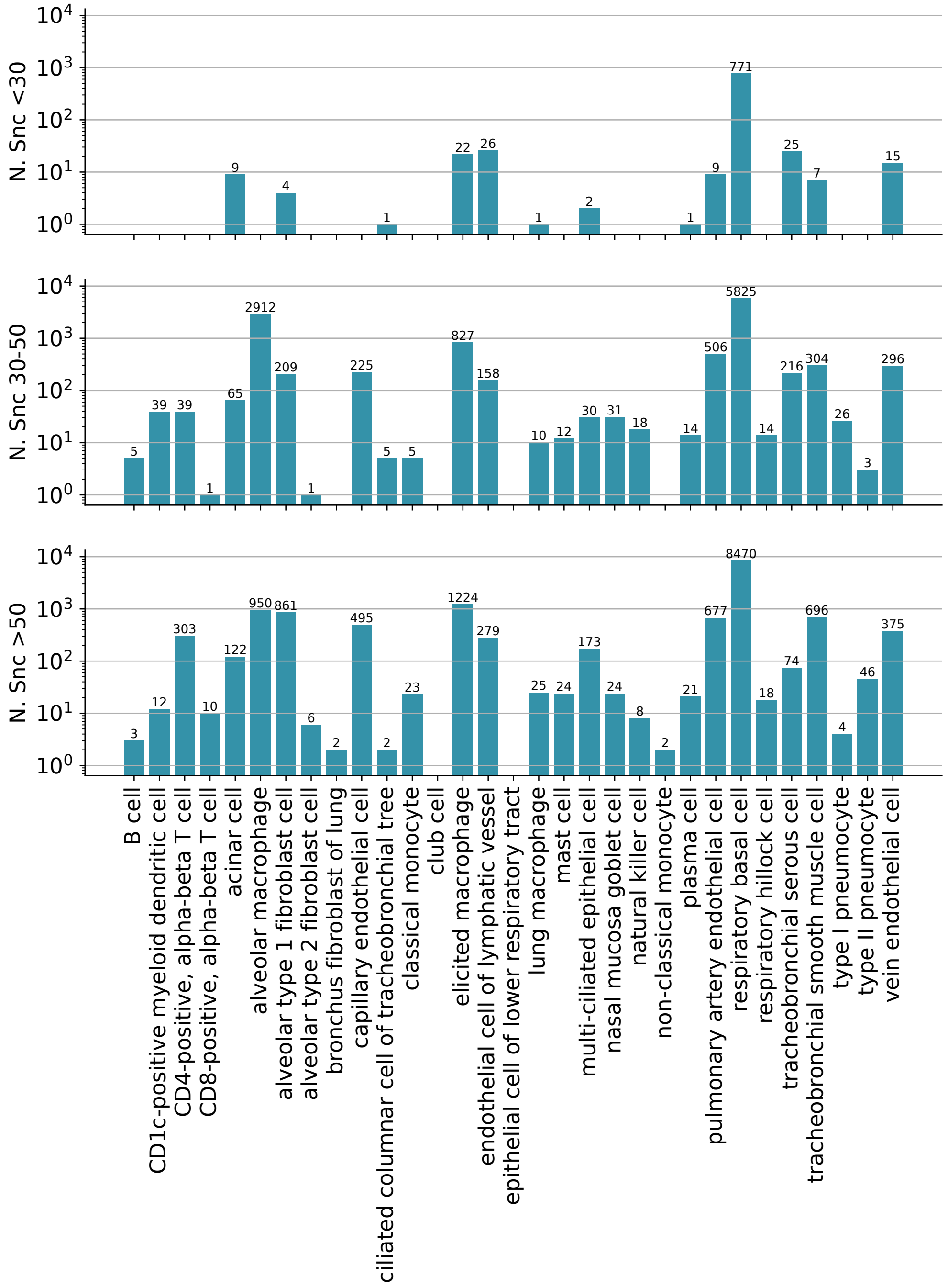


Figure 3: **Number of cells identified as senescent by the PU learner for each age group in the HLCA dataset.**

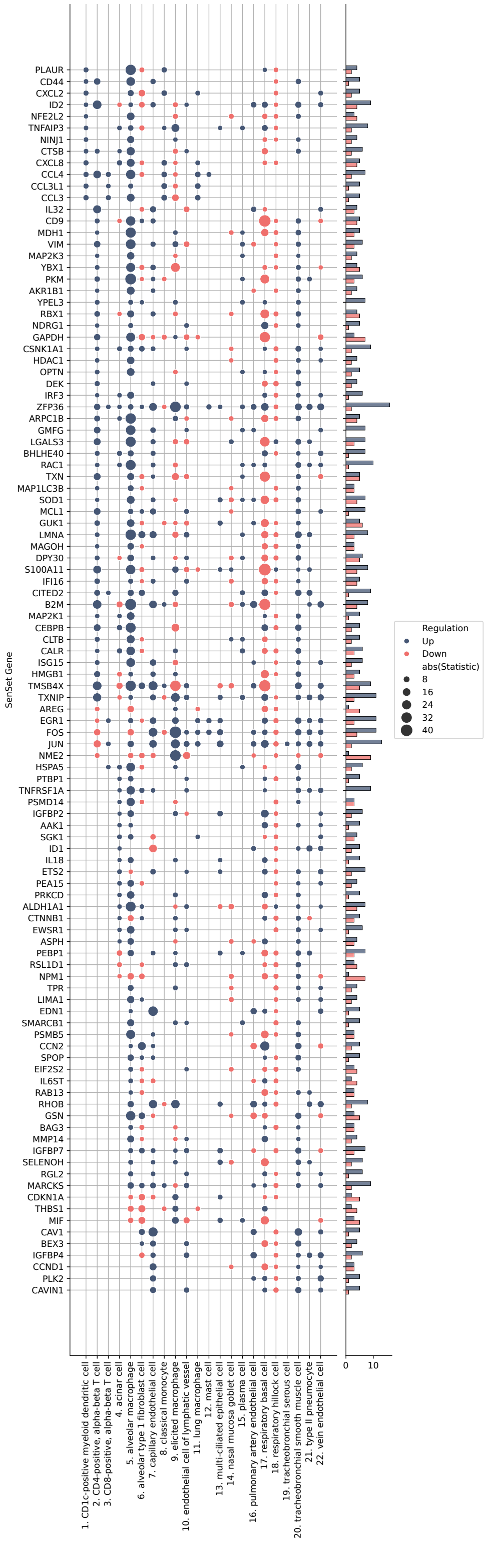


Figure 4: **All SenSet genes and their enrichment for 22 cell types in the HLCA dataset. The DE test was performed using a rank sum test.**

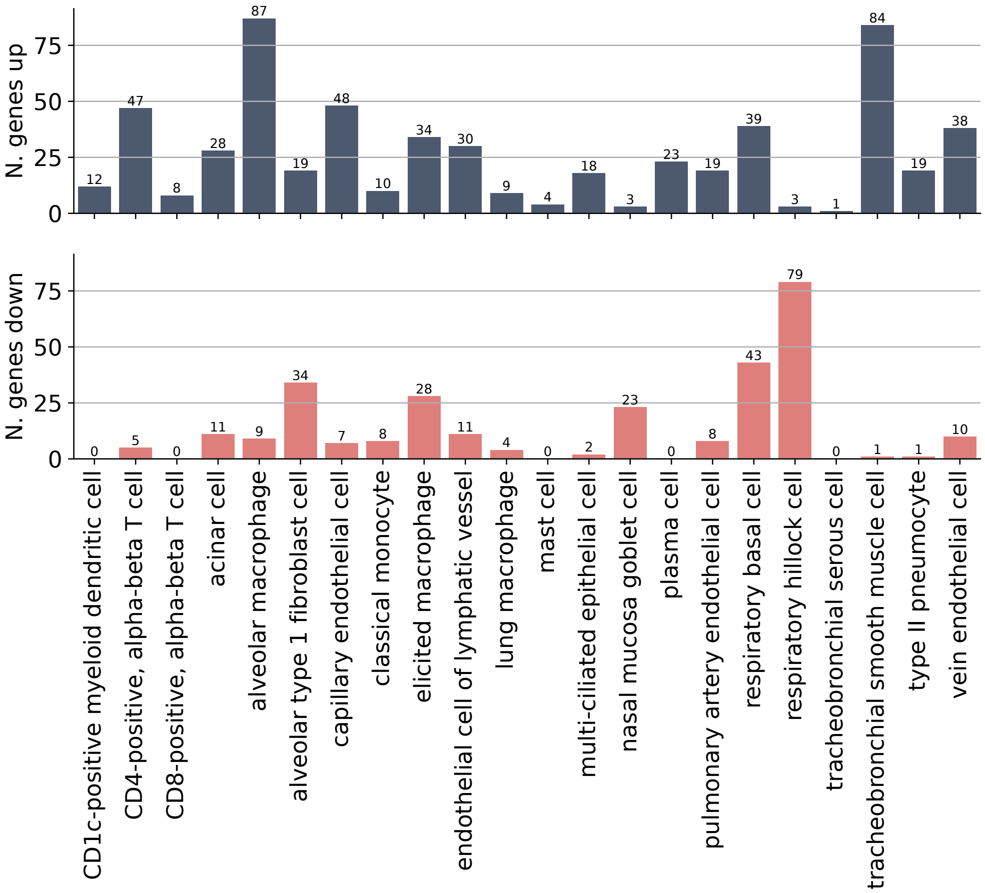


Figure 5: **Number of SenSet genes up and downregulated in the negative (senescent) class for each cell type in the oldest age group $\mathcal{A}$.**

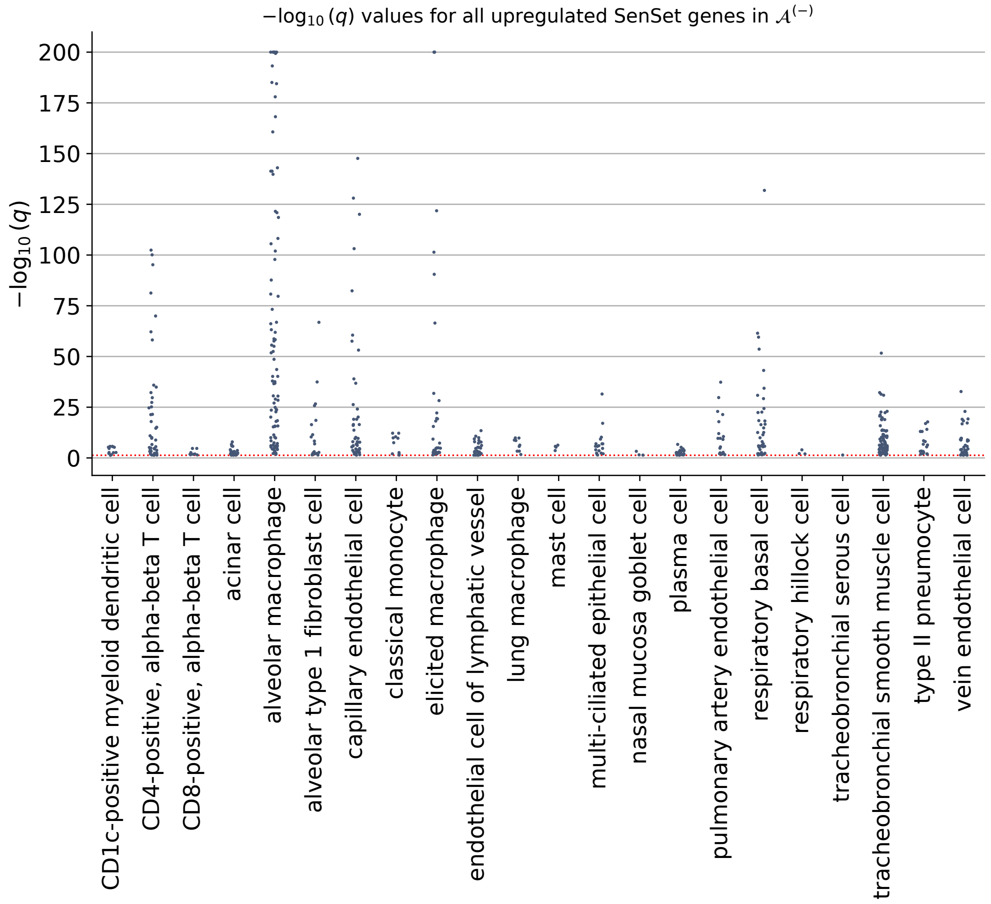


Figure 6: **Distribution of $q$-values for all SenSet genes upregulated in senescent cells across cell types in the oldest age group $\mathcal{A}$.**

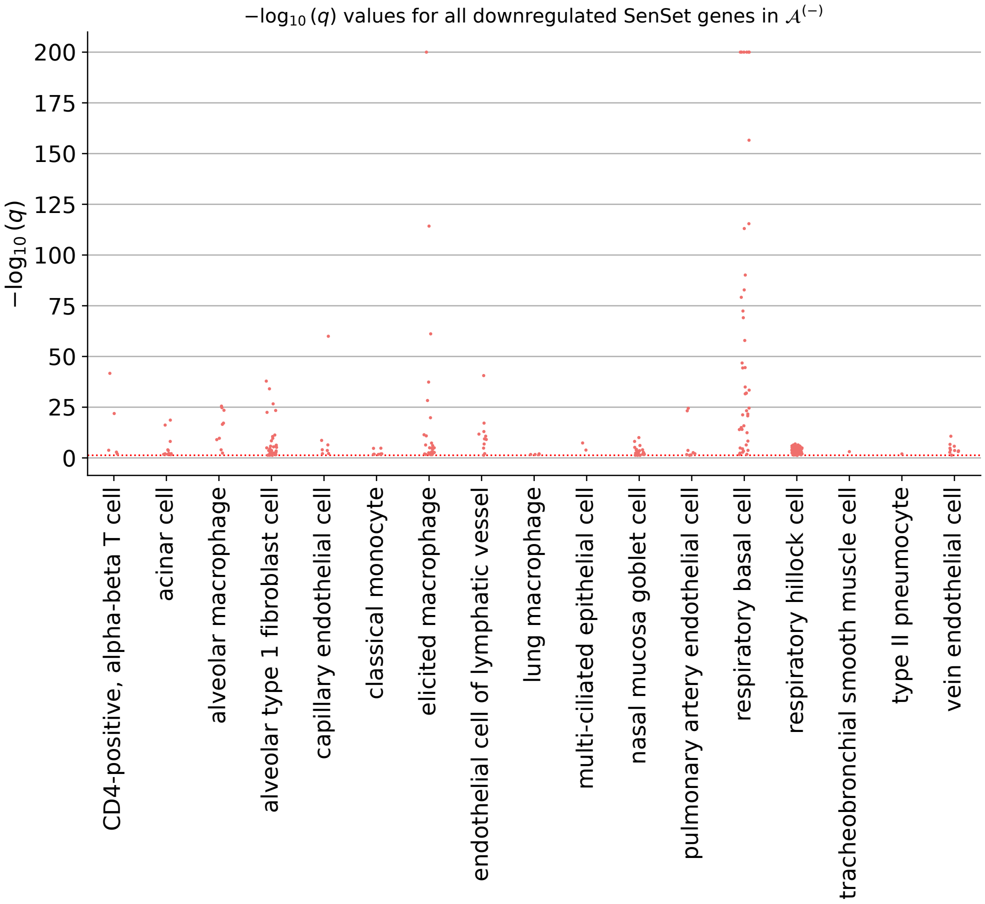


Figure 7: **Distribution of $q$-values for all SenSet genes downregulated in senescent cells across cell types in the oldest age group $\mathcal{A}$.**

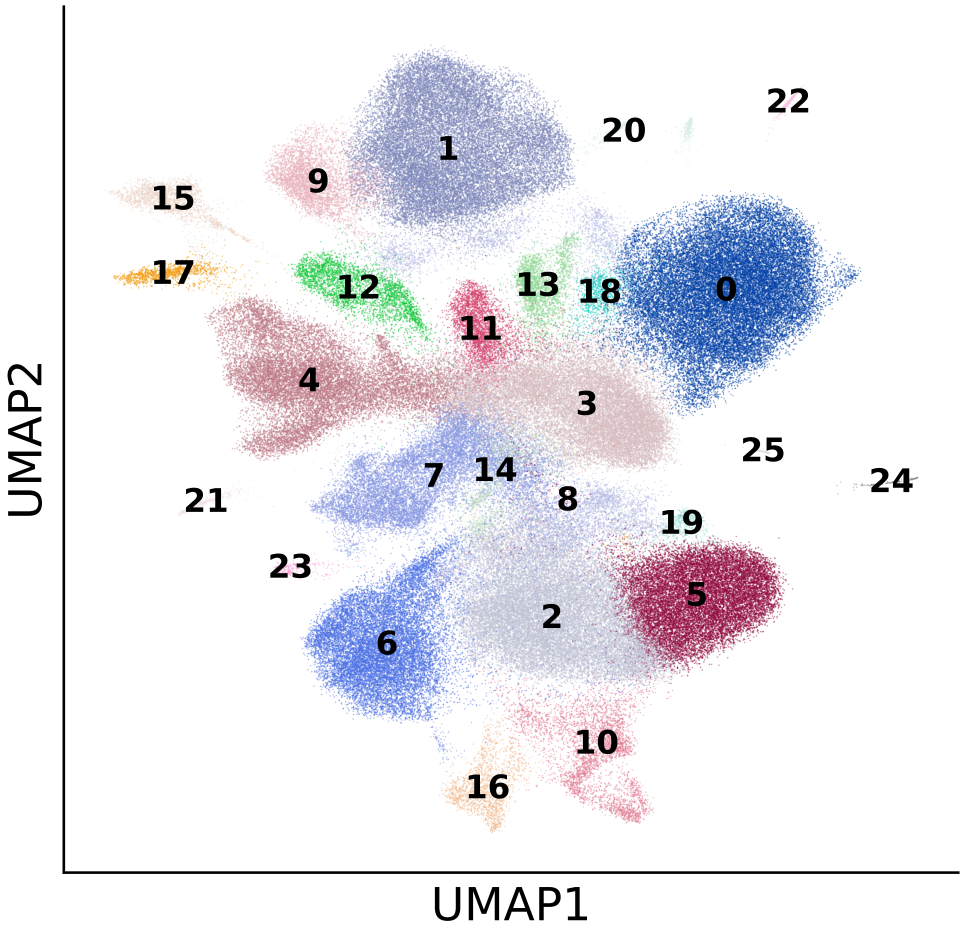


Figure 8: **All clusters identified in our PCLS data using the Leiden method.**

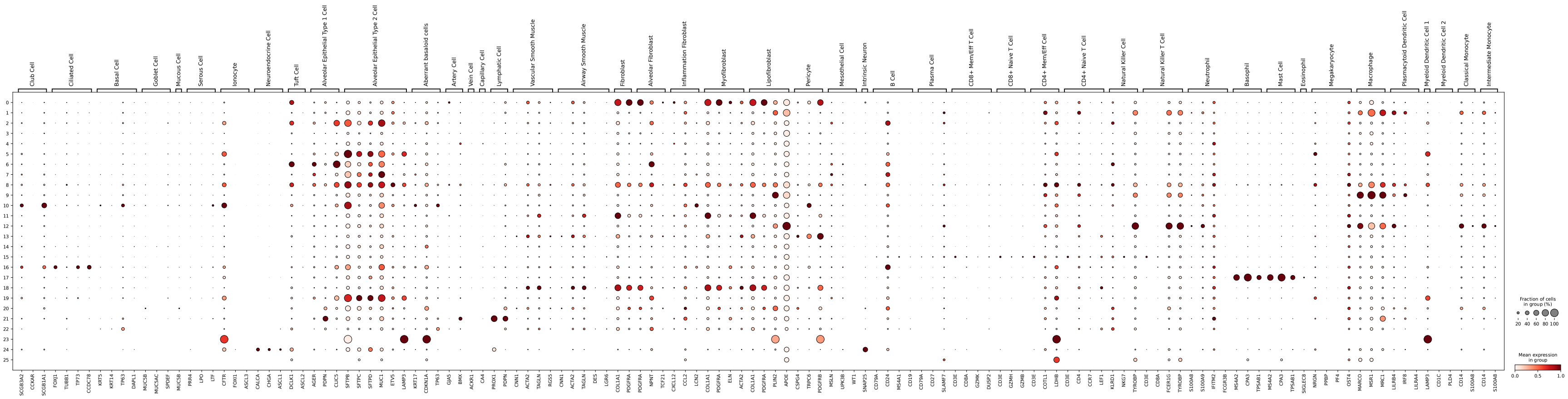


Figure 9: **Canonical markers used to identify cell types in the PCLS. These markers were obtained from Travaglini *et al.* [1] and were supplemented with additional markers for transient alveolar epithelial cells.**

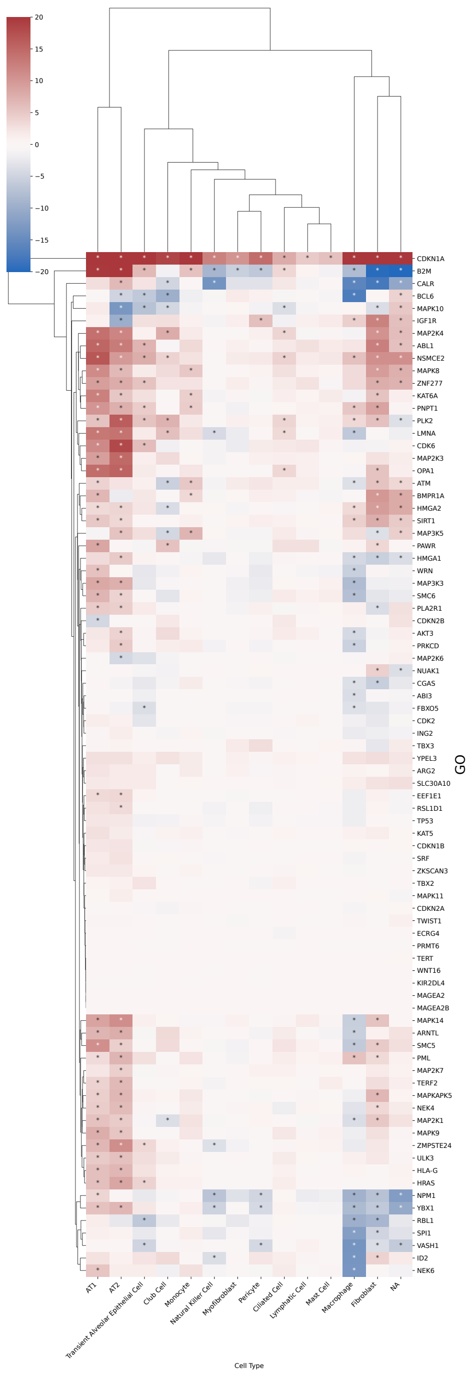


Figure 10: **Regulation of all marker genes in the GO set for each cell type in treated vs. control cells in our PCLS model.**

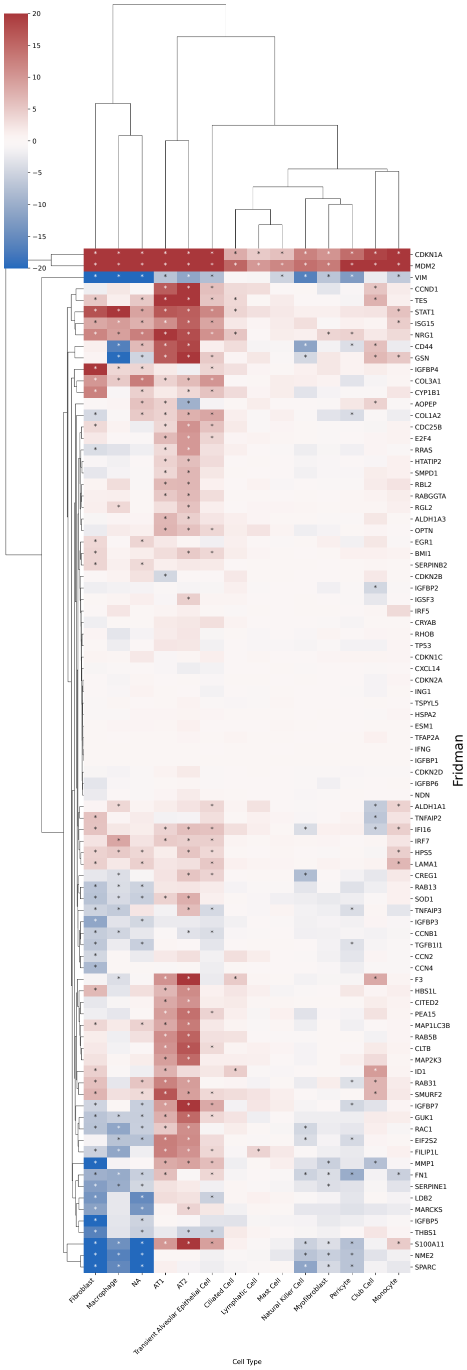


Figure 11: **Regulation of all marker genes in the Fridman set for each cell type in treated vs. control cells in our PCLS model.**

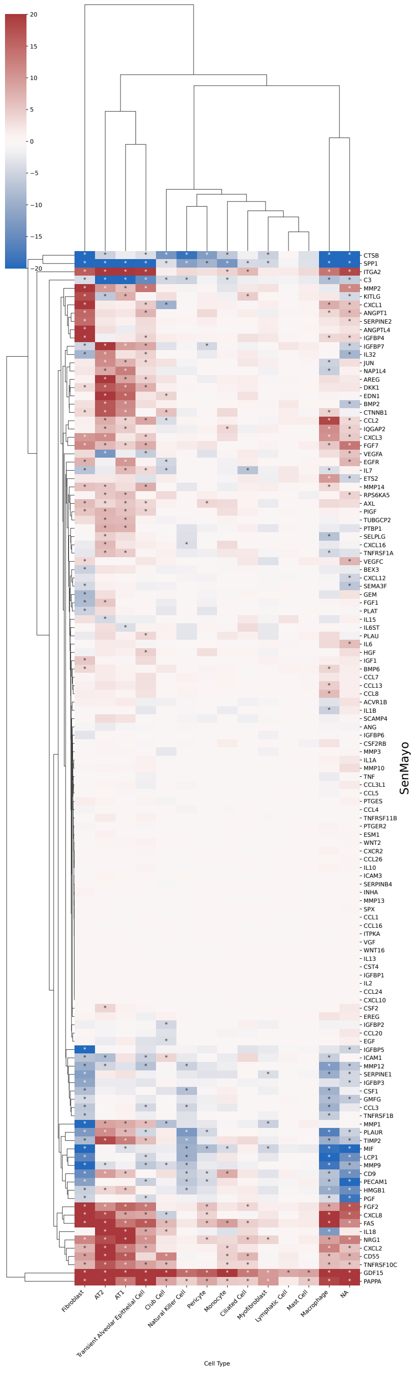


Figure 12: **Regulation of all marker genes in the SenMayo set for each cell type in treated vs. control cells in our PCLS model.**

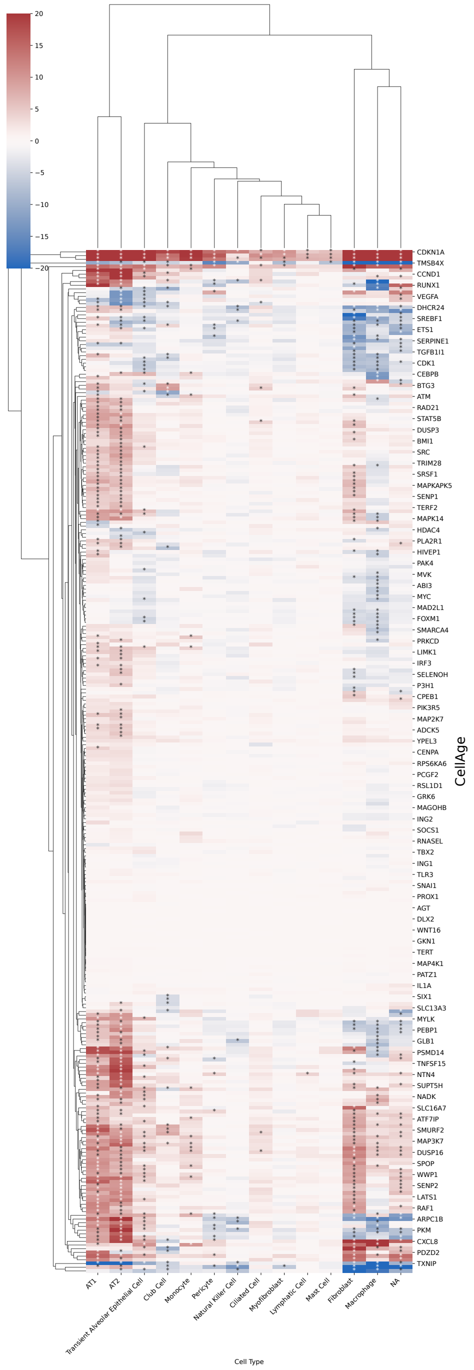


Figure 13: **Regulation of all marker genes in the CellAge set for each cell type in treated vs. control cells in our PCLS model.**
